# Network analysis of flowering time genes suggests regulatory changes among *SOC1* orthologues in response to cold in *Brassica napus*

**DOI:** 10.64898/2025.12.16.694548

**Authors:** Gurpinder Singh Sidhu, Samuel Burrows, Hugh Woolfenden, Rachel Wells, Richard J. Morris

## Abstract

Flowering plants respond to multiple environmental and endogenous cues to determine the timing of their transition from the vegetative to floral state. Most of our knowledge of the gene regulatory network (GRN) controlling flowering time has been derived from the model plant *Arabidopsis thaliana*. This knowledge needs to be translated to crop plants to support the development of varieties that can be grown in different and rapidly changing climatic conditions. However, due to increased genome complexity and limited prior knowledge, translation into crops is not always straightforward. Here, we present a study of the GRN controlling flowering time in *Brassica napus* (oilseed rape), an allotetraploid crop that is a close relative of Arabidopsis. Using a comparative transcriptomics approach, we show that the majority of the orthologous gene pairs have similar expression dynamics over development between Arabidopsis and *Brassica napus*. Some genes, however, have experienced regulatory changes, with flowering time genes in *Brassica napus* having higher than average differences in their expression profiles from their Arabidopsis orthologues. Despite these differences, the inferred GRN for flowering in *Brassica napus* exhibits a similar network topology to the network known in Arabidopsis. This is likely due to preferential retention of these genes in higher paralogue numbers, which allows subtle changes in the regulation of individual paralogues, while still conserving the overall regulatory structure through evolution. We discover and present a detailed analysis of one such example where orthologues of *SUPRESSOR OF OVEREXPRESSION OF CONSTANS 1* (*SOC1*) have similar expression patterns under normal conditions, but different dynamics under cold temperature conditions, suggesting possible subfunctionalisation among paralogues in response to temperature change.

## 1 Introduction

The switch from vegetative to reproductive growth is a key developmental transition in the life cycle of a plant. During what is known as the floral transition, the shoot apical meristem stops producing leaves and starts making flowers. Plants utilise both environmental and endogenous cues, such as ambient temperature, photoperiod, exposure to long term winter cold, age of the plant, hormones, sugars and the circadian clock, to time this transition to ensure reproductive success [4] [5]. Research using Arabidopsis mutants first paved the way to map the response to these cues to genetic loci [33] [57] [59], leading to elucidation of the regulatory framework controlling flowering time within this model plant. The multiple genetic pathways that have been characterised in Arabidopsis form a gene regulatory network (GRN) that converges on a handful of central regulators, known as floral integrator genes. These central regulators control the floral meristem identity genes to induce flowering [9] [67] [69].

Just as plants in the wild need to time their floral transition for their reproductive success, synchronised flowering at the correct time is important for maximising the agronomic outputs of crop plants. Much is known about the regulation of flowering time at the molecular level in Arabidopsis, therefore the challenge is to exploit this knowledge to achieve similar understanding in crops. Direct translation of the underlying GRN is not straightforward due to complex genomes in non-model species that, together with artificial selection under domestication, leads to differences compared to the model plant [20] [45]. For example, research in soybean (*Glycine max*) has shown that circadian clock genes controlling flowering have been the main targets of domestication. A legume specific transcription factor (TF), *E1*, along with four other orthologues of Arabidopsis flowering time genes have been identified as key components of the GRN [42] [74] [75] [79]. Orthologues of *FLOWERING LOCUS T* (*FT*) have been shown to aid adaptation in rice and maize [21] [50]. Interestingly, members of the *FLOWERING LOCUS C* (*FLC*) clade of MADS-box genes have not been reported as major flowering regulators outside of the Brassicaceae family [45].

The Brassicaceae family contains crop plants that are among the closest relatives to Arabidopsis and therefore includes prime candidates to translate research from model to crop. Brassicaceae are notable for their extensive genetic and phenotypic diversity, often within the cultivars of the same species [14]. The Arabidopsis-Brassica lineage split between 14.5 to 20.4 million years ago [81], followed by a whole genome triplication event [2]. *Brassica napus* (oilseed rape, Canola) is an allotetraploid (AACC) formed by the hybridisation of two diploid progenitors, *Brassica rapa* (AA) and *Brassica oleracea* (CC) [70]. *Brassica napus* is among the most important oil crops and grown across the globe. As flowering is a key phase for determining yield in *B. napus*, breeding efforts have aimed to time this crucial transition to maximise growth time while reducing exposure to adverse environmental conditions [61].Environmental challenges, such as drought [24] and winter warming [10], have been shown to negatively impact yield in this crop. With the predicted change in climate, most *B. napus* growing areas are set to face drought during the flowering season [41], making it more imperative than ever to gain fundamental understanding of how flowering time is regulated to underpin the development of new varieties.

Despite *B. napus* being closely related to Arabidopsis, the knowledge transfer between these species is not straightforward due to two main reasons. Firstly, since *B. napus* is cultivated across the globe, breeding efforts to select for suitable flowering time for a particular climate has produced diverse types of cultivars [38] [63]. Spring-types flower quickly and have no requirement for long-term winter cold (‘vernalisation’) to induce flowering. Semi-winter-types respond to winter cold treatment, while winter-types require a long period of winter cold before they can transition to flowering. This variation suggests that there are regulatory differences in the control of flowering time between these different types of cultivars. Secondly, the allotetraploid genome complicates the inference and analysis of GRNs. Based on its evolutionary history, *B. napus* should theoretically have six orthologues for every Arabidopsis gene, however, research has shown that there exists only a four-fold difference in the number of genes between the two species on the whole genome level [13]. This indicates gene loss over evolutionary time. Flowering time genes, however, have been shown to be preferentially retained within the genome [27] [62]. While multiple paralogues within a system can be retained immediately following duplication, it has been posited that the ultimate fate of a paralogue is to acquire deleterious mutations, becoming either non-functional (‘nonfunctionalisation’) or developing a novel function (‘neofunctionalisation’) [25] [51]. Genetic redundancy can be maintained in cases of ‘dosage-balance’ where stoichiometric ratios between gene products are necessary [6] or ‘subfunctionalisation’ where genes retain a complementary subset of functions of the ancestral gene, such that any deletions of paralogues could have negative fitness consequences [19]. Therefore, the retained orthologues of the same Arabidopsis flowering gene could have diverse expression patterns, with different dynamics over time, in different tissues or in response to different stress conditions. Indeed, there is some evidence that this is the case [12] [22] [27] [64] and therefore, it is important to study the role of each orthologue individually. As recent reviews have noted, our understanding of flowering time GRN within *B. napus* remains fragmented [45] [61] [82]

In this study, we sought to infer GRNs of orthologues of flowering time genes in *B. napus* from their expression dynamics throughout development, leading to the floral transition. We generated timeseries transcriptomic datasets of shoot apical meristems (SAMs) from two different cultivars of *B. napus*, with contrasting vernalisation requirements. By comparing the expression pattern of each gene to its Arabidopsis orthologue, we show that flowering time genes have greater differences in their expression dynamics between retained paralogues compared to the rest of the genome. Our inferred GRN in the spring-type cultivar shows that despite the divergence in dynamics, the network topology is similar to the Arabidopsis GRN. However, the network inferred for the semi-winter cultivar splits into different sub-networks or communities. Our analysis suggests this is due to cold treatment, and we discover regulatory changes between orthologues of a key floral integrator, *SUPRESSOR OF OVEREXPRESSION OF CONSTANS 1* (*SOC1*) in the semi-winter cultivar. Some orthologues of *SOC1* show rapid upregulation when plants are subjected to short-day, cold (‘winter’) conditions with their expression maxima during these conditions while others remain repressed until the floral transition. Exposing the spring-type cultivar the same short-day, cold treatment reveals that these differences in expression dynamics are independent of cultivar type. We also observe a correlated upregulation of orthologues of an upstream regulator of *SOC1*, *CIRCADIAN CLOCK ASSOCIATED 1* (*CCA1*), coupled with variations in binding site sequences in the promoter region of *SOC1* orthologues compared to Arabidopsis. While *CCA1* is a central gene of the circadian pathway, there have been reports linking its expression to cold stress in Arabidopsis [53]. Experimentally, we show that *SOC1* expression divergence and *CCA1* upregulation occur together under cold stress, independent of photoperiod change. Overall, we infer flowering time GRNs in a polyploid crop and present an analysis of similarities and subtle differences in the regulation of gene between model and crop.

## 2 Materials and Methods

### 2.1 Plant material and growth conditions

The data presented in this study is collected from two *B. napus* cultivars, Stellar and Zhongshuang 11 (ZS11). Stellar is a spring-type cultivar that does not require vernalisation to undergo floral transition while ZS11 is a Chinese semi-winter-type cultivar that responds to vernalisation prior to floral transition. Plants were sown in cereal mix (40 % medium grade peat, 40 % sterilised soil, 20 % horticultural grit, 1.2 kg/m^3^ PG mix 14–16–18 + Te base fertilizer, 3 kg/m^3^ maglime, 300 g/m^3^ Exemptor). Following germination, each seedling was transplanted to a 5 cm *×* 5 cm *×* 4.5 cm cell in a standard 24 cell-tray. Plants were grown in controlled environments with a 16-hour photoperiod. Temperatures were set to 18 *^◦^*C during the day and 15 *^◦^*C during the night, with relative humidity maintained at 70 %. Short-day, cold conditions within this study refer to an 8-hour photoperiod at constant 5 *^◦^*C temperature.

For the experiment shown in Figure 1, plants were grown in a Conviron MTPS 144 controlled environment room with Valoya NS1 LED lighting (250 *µ*mol/m^2^s). Stellar plants were grown under normal conditions throughout with floral transition, marked by visible floral primordia at the shoot apical meristem (Shown in Supplemental Figure S6), occurring at day 30. ZS11 plants were grown in normal conditions but at day 21, were moved to short-day, cold conditions. They were transferred back to normal conditions after three weeks (day 42). For experiments shown in Figure 6 (a) and Figure 8, plants were grown in Hettich 1700 growth cabinets with 200 *µ*mol/m^2^s lighting. Stellar plants were grown under normal conditions and transferred at day 21 to one of three different treatments: (1) short-day, cold conditions (Figure 6 (a)), (2) short-day conditions, or (3) cold conditions (Figure 8). The plants were transferred back to normal conditions after two weeks (day 35).

**Figure 1:**
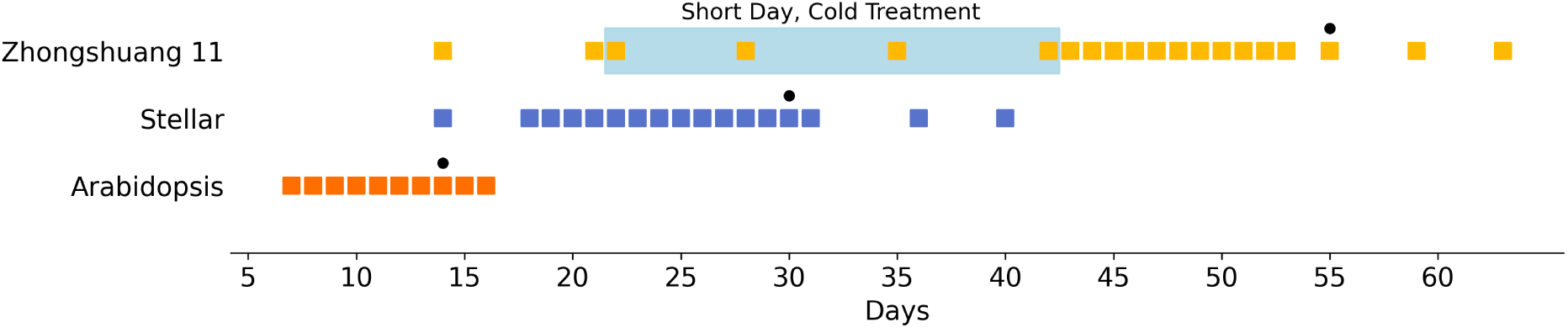
Sample timepoints for *B. napus* cultivars Stellar, Zhongshuang 11 (ZS11) and Arabidopsis (Col-0). The square markers show timepoints when shoot apical meristems (SAMs) were sampled for RNA-seq for the spring-type *B. napus* cultivar Stellar and the semi-winter cultivar ZS11. ZS11 plants were given three weeks of shortday, cold (‘winter’) treatment prior to the floral transition, shown in the figure as a light blue bar. Similar data was obtained for Arabidopsis (Col-0) from Klepikova *et al*. [32]. SAMs were monitored under the microscope and floral transition times were recorded, marked as ‘•’ in the figure. See Supplemental Figure S6 for images of vegetative and floral SAMs. Supplementary Tables 1 and 2 show the details of the sampled timeseries, alongwith alignment statistics.

### 2.2 Sampling and RNA sequencing

Three replicates, each comprising a pool of three dissected shoot apical meristems were collected for every timepoint in the timeseries presented in this study. Following dissection, samples were immediately frozen in liquid nitrogen. Samples were ground to a fine powder for RNA-extraction using the EZNA ® Plant RNA Kit (Omega Bio-tek Inc) following the manufacturer’s instructions. RNA samples were processed by Novogene, with paired end 150 bp library preparation using NEB next ultra directional library kit (New England Biolabs) to generate a minimum of 7.5 G raw data per sample. Each timepoint shown in Figures 1, 6 (a) and 8 thus resulted in 3 RNA-seq samples following QC, except for timepoint 53 in ZS11 timeseries shown in Figure 1, which only has one replicate.

### 2.3 RNA-seq data processing

FastQC (version 0.11.9) [1] and MultiQC (version 1.29) [18] were used for QC and data collation respectively. Trimmomatic (version 0.39) [7] was used to remove any adaptor contamination with the following flags: ILLUMINACLIP:TruSeq3-PE-2.fa:2:30:10:1:true, HEADCROP:15, SLIDINGWINDOW:4:15 and MINLEN: 50. HISAT2 (version 2.1.0) [30] was used to align the data to Darmor v10 reference genome [60] with ‘RF strandedness’. Samtools (version 1.9) [16] was used for sorting and indexing. The resultant bam files were filtered to only keep uniquely mapped reads. StringTie (version 2.1.1) [55] was used for expression quantification, with TPM (‘Transcripts Per Million’) [71] selected as the measure of gene expression. Supplementary Tables 1, 2, 5 and 8 show detailed alignment statistics for each timeseries presented. ‘_BnaDAR’ suffix has been omitted from Darmor v10 gene names mentioned throughout this manuscript.

### 2.4 Orthologue mappings and determination of flowering time genes

A reciprocal BLAST based strategy was used to determine one-to-many orthologue mappings between the Arabidopsis TAIR10 genome [36] and the *B. napus* Darmor v10 reference [60]. For an Arabidopsis gene, if any hits (e-value *<* 1e-5) within the *B. napus* genome also had the same Arabidopsis gene as the best hit in reciprocal blast search, they were marked as a putative orthologue pair. The custom python script used to perform this search is available on zenodo: https://doi.org/10.5281/zenodo.17631422. The set of 306 Arabidopsis genes involved in flowering time regulation were obtained from the FLOR-ID database [9]. The reciprocal BLAST search identified 1,232 orthologues of these flowering time genes in *B. napus*.

### 2.5 Curve registration

greatR (version 2.0.0.9000) [34] was used to perform curve registration. For both Stellar and ZS11 registrations against the Col-0 timeseries, Col-0 was used as the query accession. ‘Z-score’ scaling method was used and overlapping percentage between the curves was set to 75. The rest of the parameters were set to their default values. greatR filters low expressed genes by default. 1071 genes in Stellar and 1069 genes in ZS11 passed this filtering criteria and were used for curve registration analysis. Full set of results are available in Supplemental File 1.

### 2.6 Network inference and analysis

To create a high confidence list of transcription factors (TFs) from the *B. napus* Darmor v10 reference, we identified the orthologues of Arabidopsis TFs from PlantTFDB database [26]. Protein sequences for these genes were used as an input for DeepTFactor tool [31]. The TF orthologues that were also classified as TFs by DeepTFactor were included in the list of high confidence TFs. Orthologues of flowering time genes that had an average TPM value across timeseries greater than 0.001 and at least 30 % of the timepoints with TPM *>* 0 were used for network inference. Out of 1,232 orthologues of flowering time genes, 980 and 976 genes passed these criteria in Stellar and ZS11. Out-predict [15] was used for network inference with default parameters. The input data and outputs for all inferred networks are available in Supplemental File 2. Inferred networks were filtered to remove self-loops and edges with importance scores less than 0.001 to remove likely false positive edges. This resulted in networks with 16,791 and 20,314 edges for Stellar and ZS11 timeseries respectively. To create ‘high confidence’ networks for visualisation purposes, we selected the top 500 edges based on importance scores, resulting in networks with 304 and 377 nodes for Stellar and ZS11 respectively. NetworkX (version 3.1) [23] was used for visualisation (Spring layout; seed 123 for Figure 3), degree centrality, average clustering coefficient analysis, community detection using fluid communities algorithm [52] and modularity calculation for the identified communities. A seed of 246 was used for community analysis.

### 2.7 Multiple sequence alignment

For promoter sequence alignment, sequences up to 2000 bp upstream of the start codon were isolated from TAIR 10 [36] and Darmor v10 [60] genome assembles. Protein sequences were also obtained from the same assemblies. The MAFFT online server [29] was used to perform the alignment in both cases using the FFT-NS-i method, allowing automatic inference of sequence direction to allow best possible alignments. Jalview (version 2.11.5.0) [76] was used for visualisation and calculation of pairwise identities.

## 3 Results

### 3.1 Orthologues of Arabidopsis flowering time genes in *B. napus* show more differences in their expression dynamics than the rest of the genome

Flowering time genes have been shown to be preferentially retained in the *B. napus* genome [27]. This state of retained redundancy provides more possibilities for change in regulation that can lead to paralogues diverging in their expression, likely aiding in adaptation to new conditions or acquiring new functions [44]. As an example, previous research into the orthologues of *FLC* gene in *B. napus* has shown that while the summed expression of orthologues correlates with vernalisation requirements of different cultivars, individual paralogues show different expression dynamics in response to vernalisation conditions [12]. To test whether this is a widespread occurrence in the genome, or specific among preferentially retained flowering time genes, we selected two cultivars of *B. napus*, Stellar and Zhongshuang 11 (ZS11), with contrasting vernalisation requirements to gather data on expression dynamics throughout plant development. Stellar is a spring-type cultivar that does not require vernalisation to undergo floral transition, while ZS11 is a Chinese semi-winter cultivar that responds to vernalisation. Figure 1 shows the timepoints when shoot apical meristem tissue was sampled for RNA-seq for the two cultivars from day 14 until the plants reached BBCH51 developmental stage [8]. Stellar plants underwent the floral transition at day 30, while ZS11 plants floral transitioned on day 55 after three weeks of vernalisation treatment from day 21 to day 42.

We sought to compare the time series expression data obtained from this experiment against a publicly available similar dataset obtained from *A. thaliana* Col-0 cultivar [32]. If paralogues have conserved expression patterns, their expression dynamics should follow a similar curve to their Arabidopsis orthologue. To facilitate the comparison between different developmental times across species, we used curve registration [58] [11]. Supplemental Figure S1 shows that curve registration is indeed required for a comparison of time series transcriptomic data across species growing on different timescales. Using this technique, we can also reproduce the findings of previous reports and show that individual *FLC* orthologues have diverged in their expression dynamics (Supplemental Figure S2).

Extending the curve registration analysis to all expressed genes shows that for both cultivars, the majority of genes show similar dynamics to their Arabidopsis orthologues, with 60,438 genes out of 72,239 expressed genes in Stellar and 55,332 genes out of 71,382 expressed genes in ZS11 registered to their corresponding Arabidopsis orthologue (Figure 2 (a)). This suggests that most genes have conserved expression dynamics, and hence, could be performing similar functions in the two species. The proportion of genes with conserved dynamics is slightly lower in ZS11 (78 % vs 84 %), which could be because the plants were given winter cold treatment, while the Col-0 plants were not. This environmental variation may have resulted in changes in gene expression that would have not occurred in Col-0 and Stellar in the absence of that treatment. For both cultivars, however, this proportion of registered genes drops among the subset of orthologues of flowering time genes. Only 887 out of the 1103 expressed orthologues to flowering time genes in Stellar (80 %) and 789 out of 1100 in ZS11 (72 %) were registered.

**Figure 2:**
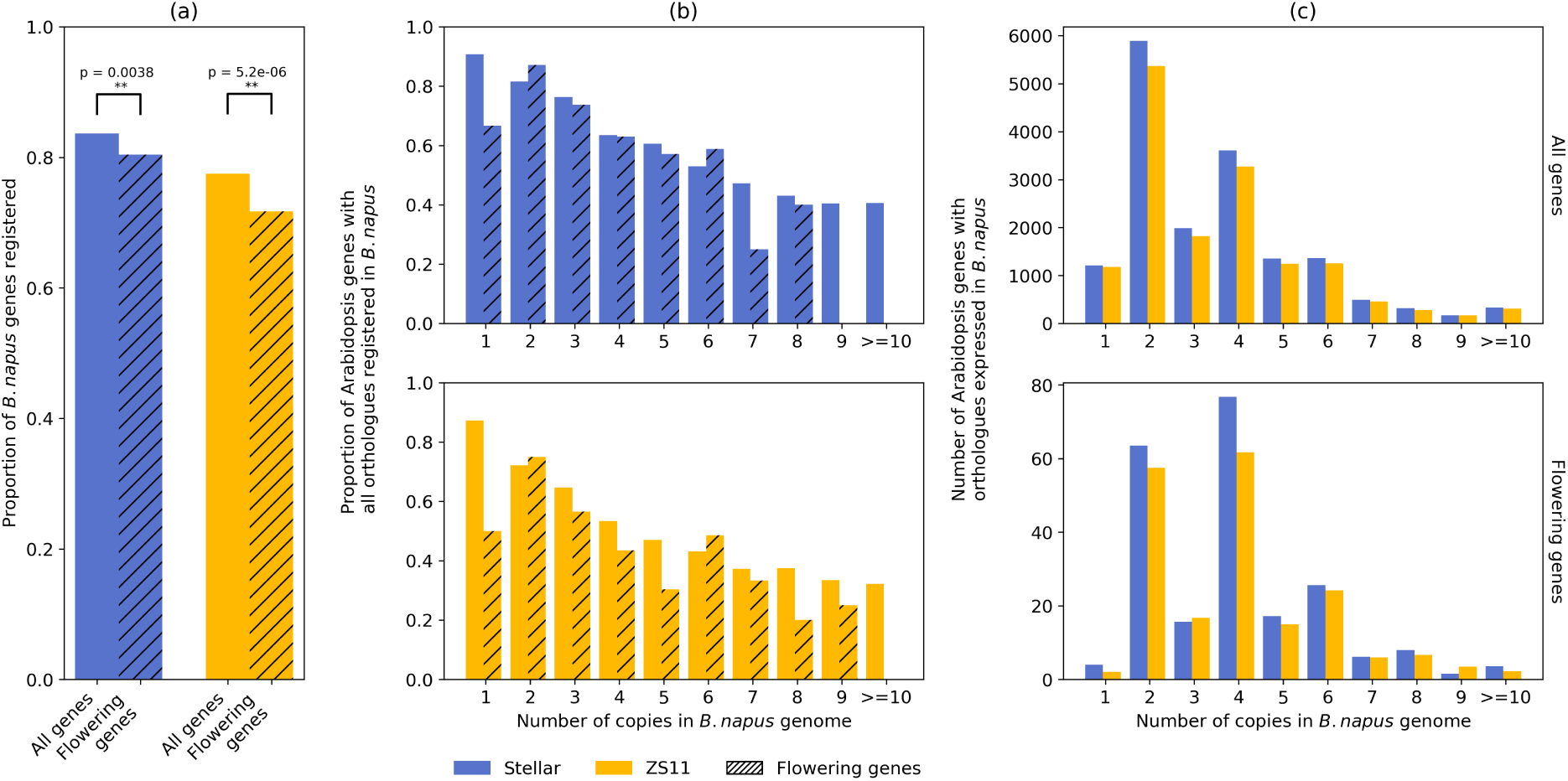
Orthologues of flowering time genes show more varied expression dynamics and are preferentially retained in higher paralogue numbers compared to rest of the genome. We used curve registration to compare expression dynamics over time in shoot apical meristem (SAM) between Arabidopsis (Col-0) and *B. napus* cultivars (Stellar and ZS11). (a) In both cultivars, the proportion of genes with the same expression dynamics as their Arabidopsis orthologue is significantly lower for flowering time genes compared to the whole genome (** p-value < 0.05; proportions z-test; using statsmodels python package [66]). Shown in Supplemental Figure S3, we also conducted bootstrapping-based analysis to show that this divergence for the set of flowering time genes cannot occur by random chance. (b) The propensity for expression dynamics to diverge is higher among genes retained in higher number of paralogues or copies within *B. napus* genome. (c) We find that orthologues of flowering time genes are preferentially retained and expressed within our dataset, correlating with their higher degree of divergence compared to the rest of the genome. For both cultivars, the most common number of expressed paralogues for the whole genome is two, while for flowering time genes, this number increases to four.

To show that this drop is not reproducible by sampling a random set of genes from the genome we performed a bootstrapping analysis. Flowering time genes were replaced with samples of equal size (1103 for Stellar and 1100 for ZS11) from the set of all genes 10,000 times (Supplemental Figure S3). The proportion of flowering time genes registered did not fall within the distributions obtained from this analysis, demonstrating that the orthologues of flowering time genes indeed exhibit significant differences in expression dynamics compared to the rest of the genome. The full results from the curve registration analysis are in Supplemental File 1.

The preferential retention of orthologues of flowering time genes is correlated with this observed higher degree of divergence. For both cultivars, as the number of orthologues in *B. napus* increases, the proportion of *B. napus* orthologues with all the copies registered to the corresponding Arabidopsis gene decreases (Figure 2 (b)). Furthermore, we also observe that flowering time genes are expressed in higher copy numbers compared to the set of all genes, thus corroborating previous studies [27] [62]. The most common number of paralogues in the genome for all genes is two, while for the set of flowering time genes, this increases to four (Figure 2 (c)). Hence, on a whole genome level, as the number of retained paralogues increases, the propensity for accumulating regulatory changes increases and, since flowering time genes are preferentially retained and expressed in higher numbers of paralogues, we observe a higher relative proportion of different expression dynamics among flowering time paralogues.

### 3.2 The GRN controlling flowering time is similar between *B. napus* and Arabidopsis, however short-day, cold treatment induces changes in network structure

The GRN controlling flowering time in Arabidopsis consists of multiple pathways that integrate different environmental and endogenous cues. These pathways converge on a set of ‘floral integrator genes’ that govern the transition to flowering in the shoot apical meristem [4] [67] [69]. Given that flowering time genes are preferentially retained and exhibit differences in their expression dynamics in B. napus, we sought to determine whether the overall flowering time GRN might be similar to the one known in Arabidopsis.

To investigate network structure, we inferred the Gene Regulatory Networks (GRNs) controlling flowering time in both cultivars of *B. napus*. The timeseries data captures the expression dynamics of every gene in Stellar and ZS11 and can be used to infer probable regulatory interactions between genes. The problem of reverse engineering a GRN from expression data is mathematically challenging, and prone to errors [46] [54]. Keeping these limitations in mind, we filtered our dataset to only include orthologues of genes already known to be involved in the flowering time GRN in Arabidopsis and limited our inferences to general trends observable from the network structures. We provide the inputs, final outputs and edge weights for the inferred GRNs in Supplementary File 2.

Network inference was performed on a set of 980 and 976 genes for Stellar and ZS11, respectively. This resulted in networks with 16,791 and 20,314 edges following initial filtering (see Methods). As these networks likely contained several false positive edges and were too large for visualisation, we filtered to only include the top 500 edges based on importance scores to create ‘high confidence networks’. Figure 3 (a) and (b) show the largest interconnected component in these networks for Stellar and ZS11 cultivars. The sizes of the nodes are proportional to their degree centrality, i.e. nodes connected with a higher number of edges are larger. The top 10 nodes in both networks are labelled along with the illustration of the pathways in which their Arabidopsis orthologues are known to be involved.

**Figure 3:**
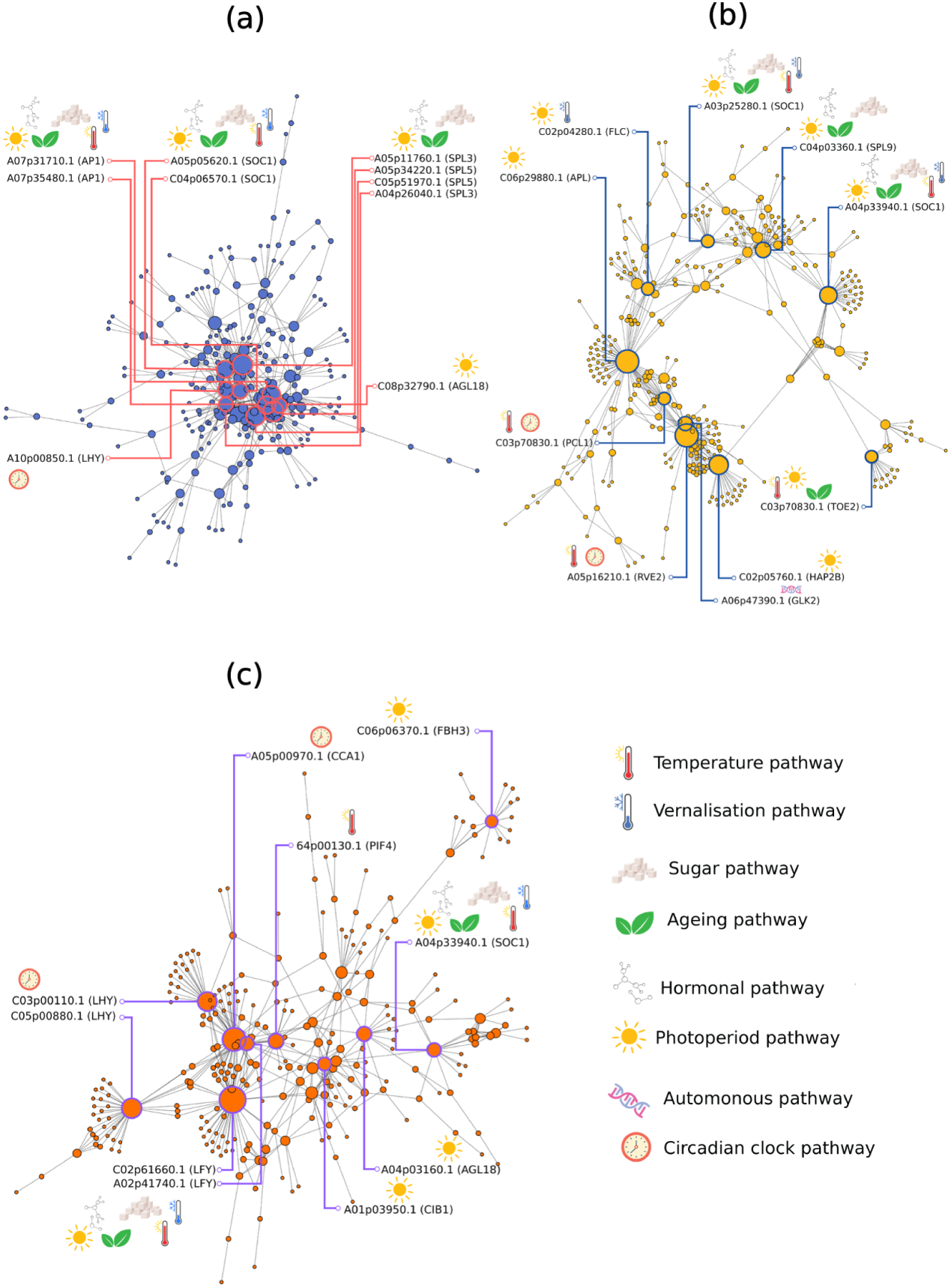
The inferred gene regulatory network (GRN) in *B. napus* follows a similar structure to Arabidopsis, with changes induced under short-day, cold treatment. We inferred networks to study regulatory relationships between flowering time gene orthologues. These network plots show the largest interconnected components among the top 500 edges, based on importance scores. The nodes represent genes, with edges showing probable regulatory relationships. The size of the nodes represents their degree centrality, with higher connected central nodes appearing larger. The top 10 nodes based on degree centrality in each network are labelled, along with the pathways inferred from their Arabidopsis orthologues. (a) The Stellar GRN follows a single community structure, with genes that are orthologues of known floral integrators appearing as key nodes in the network. (b) The ZS11 GRN lacks the single community structure visible in Stellar. Instead the network appears to comprise local clusters or communities. (c) The inferred GRN from the ZS11 timeseries using only post vernalisation data reverts to a structure similar to the Stellar network shown in (a), indicating that expression changes induced by short-day, cold vernalisation treatment, may lead to the changes in GRN structure between Stellar and ZS11.

It is clear from the structure of the inferred network from Stellar timeseries that it converges on a few key nodes, visualised in the centre with high degree centrality, including orthologues of genes from *SQUAMOSA PROMOTER BINDING PROTEIN-LIKE* (*SPL*) family, *SPL3* and *SPL5*. n Arabidopsis, these genes are up-regulated during photoperiod-induced flowering [65] and are regulated by microRNAs, specifically miRNA-156, which is involved in the ageing pathway [72] [78]. These genes have also been shown to be part of a module with *SUPRESSOR OF OVEREXPRESSION OF CONSTANS 1* (*SOC1*) that incorporates signals from gibberellic acid, the endogenous hormonal pathway [28]. *SPL* genes encode for a transcription factor that regulates floral integrator genes, *APETALA 1* (*AP1*), *LEAFY* (*LFY*) and *AGAMOUS-LIKE 8* (*AGL8*) or *FRUITFUL* (*FUL*) [80]. The *SPL* genes are present alongside orthologues of floral integrators *SOC1*, *AP1* and *AGL8* as the most important nodes in the network. Supplementary table 3 contains the extended list of top 25 nodes from the network.

The inferred flowering time network in *B. napus* cv. Stellar maintains a similar overall structure to the Arabidopsis network with the orthologues of floral integrator genes identified as nodes with high degree centrality. Exceptions are the orthologues of the genes *LATE ELONGATED HYPOCOTYL* (*LHY*) and *AGAMOUS-LIKE* (*AGL18*), which are involved in circadian clock and photoperiod pathway respectively. These genes are present as top nodes but have not been shown to integrate signals from multiple flowering pathways in Arabidopsis.

However, the network topology changed for ZS11, the semi-winter cultivar that was given 3 weeks long short-day, cold treatment. The inferred network splits into distinct subnetworks rather than converging on a set of floral integrator genes (Figure 3 (b)). The average clustering coefficient for the ZS11 network is 0.05, which is lower than 0.11 for the network inferred from Stellar data, further suggesting a dispersed network structure. We speculated that this effect is due to the short-day, cold treatment as a part of the network consists of orthologues of genes known be involved in the photoperiod pathway, as well as *FLOWERING LOCUS C* (*FLC*), a gene involved in the vernalisation pathway in Arabidopsis. The rest of the network contains floral integrators as important nodes, alongside orthologues of *PHYTOCHROME INTERACTING FACTOR 4* (*PIF4*). *PIF4* binds to *FT* and act as part of the ambient temperature pathway [35]. Supplementary table 4 contains the extended list of top 25 nodes from the network.

To test whether short-day, cold treatment causes the change in GRN structure in ZS11 compared to Stellar, we performed network inference using only data from timepoints after the treatment. The resulting inferred network, shown in Figure 3 (c), has a structure resembling that of Stellar, Figure 3 (a), and the *FLC* dominated subnetwork is no longer visible, while other floral integrators still are present among the most important nodes. We find that, despite differences in flowering time gene expression dynamics, the inferred network in *B. napus* shares similarities with the GRN structure in Arabidopsis. Orthologues of floral integrators play a key role in flowering time control, highlighting conserved regulation between *B. napus* and Arabidopsis. However, different environmental conditions result in changes to the structure of the inferred GRN, requiring further investigation.

### 3.3 Orthologues of *SOC1* in *B. napus* cv. ZS11 exhibit different expression dynamics in response to short-day, cold treatment

As floral integrator genes and their orthologues are key components of the flowering time GRNs in Arabidopsis and *B. napus* cv. Stellar, we sought to determine their position in the inferred network from *B. napus* cv ZS11.

We used a community detection algorithm, fluid communities [52], to determine clusters of nodes in the full ZS11 network (976 genes; 20,314 edges). Based on the maximum modularity metric [49] (Supplemental Figure S4), the network was divided into three communities comprising 374, 341 and 261 genes. Figure 4 shows the scaled expression profiles of the genes in these three communities, with expression profiles of individual orthologues for each floral integrator highlighted against the background of all genes in that community.

**Figure 4:**
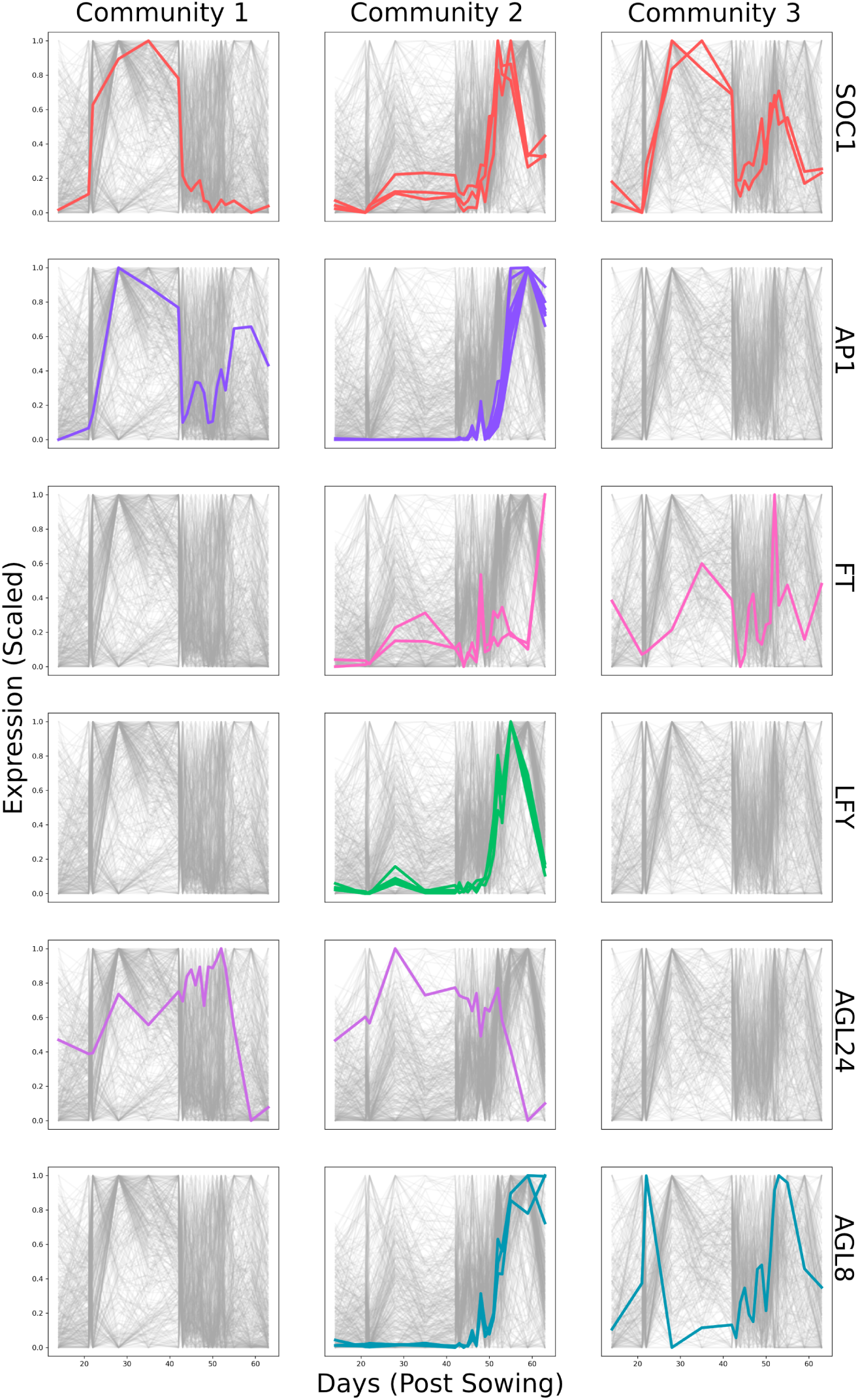
Orthologues of *SOC1* are unique among floral integrators in occupying all three identified communities within the ZS11 network. We used community detection algorithms to identify and divide the ZS11 network into three communities containing 374, 341 and 261 genes respectively. Scaled expression profiles of all the genes occupying those communities are shown in grey. In each row, the expression profile of all orthologues of a floral integrator are highlighted within each of the communities. Most orthologues occupy community 2 with only one orthologue each of *AP1* and *AGL24* in community 1 and one orthologue each of *FT* and *AGL8* in community 3. However, the six *SOC1* orthologues are spread across all three communities within the network, with one, three and two orthologues in communities 1, 2 and 3 respectively. As evident from the expression profiles, the orthologues have variations in their dynamics, indicative of divergence in regulation.

Most orthologues of the floral integrators are present in community 2, where where gene expression changes following short-day, cold treatment. Communities 1 and 3 consist of genes that undergo expression changes in response to short-day, cold treatment. Orthologues of floral integrators tend to follow similar expression dynamics, resulting in them being in one community, with at most one orthologue in another. However, orthologues of *SOC1* do not follow this trend.

The six *SOC1* orthologues are distributed in all three communities, demonstrating the high diversity in their expression dynamics. The three orthologues present in communities 1 and 3 show substantial upregulation when the plants undergo vernalisation treatment, with their maximum expression occurring during this period. Orthologues of *SOC1* are the only floral integrators that are present in all three identified communities in the network inferred from ZS11 data.

The expression patterns of the six orthologues over time in both cultivars is shown in Figure 5. All six follow similar expression dynamics in Stellar, where their expression is minimum at the beginning of the timeseries, increasing as the plant develops, reaching a maximum around the floral transition and declining thereafter. In ZS11, these genes have different expression profiles. A03p25280.1 (*Bna.SOC1.A03*) shows the least change in expression as plants are exposed to short day, cold conditions. A05p05620.1 (*Bna.SOC1.A05*) shows some upregulation, while the C genome copies, C03p30160.1 (*Bna.SOC1.C03*), C04p06570.1 (*Bna.SOC1.C04a*) and C04p71550.1 (*Bna.SOC1.C04b*) have their maximum expression during the vernalisation period. Following vernalisation, only *Bna.SOC1.A03* and *Bna.SOC1.A05* show upregulated expression leading up to floral transition. *Bna.SOC1.C03* and *Bna.SOC1.C04a* also increase their expression levels, however, to a lesser extent than when plants were undergoing vernalisation. These expression differences lead to differences in inferred regulators among *SOC1* orthologues between the two GRNs discussed earlier. To compare the sets of potential upstream regulators, we performed Rank Biased Overlap [77] analysis and found that *SOC1* orthologues are unique among the floral integrators, suggesting major differences in the regulation of paralogues (Supplemental Figure S5).

**Figure 5:**
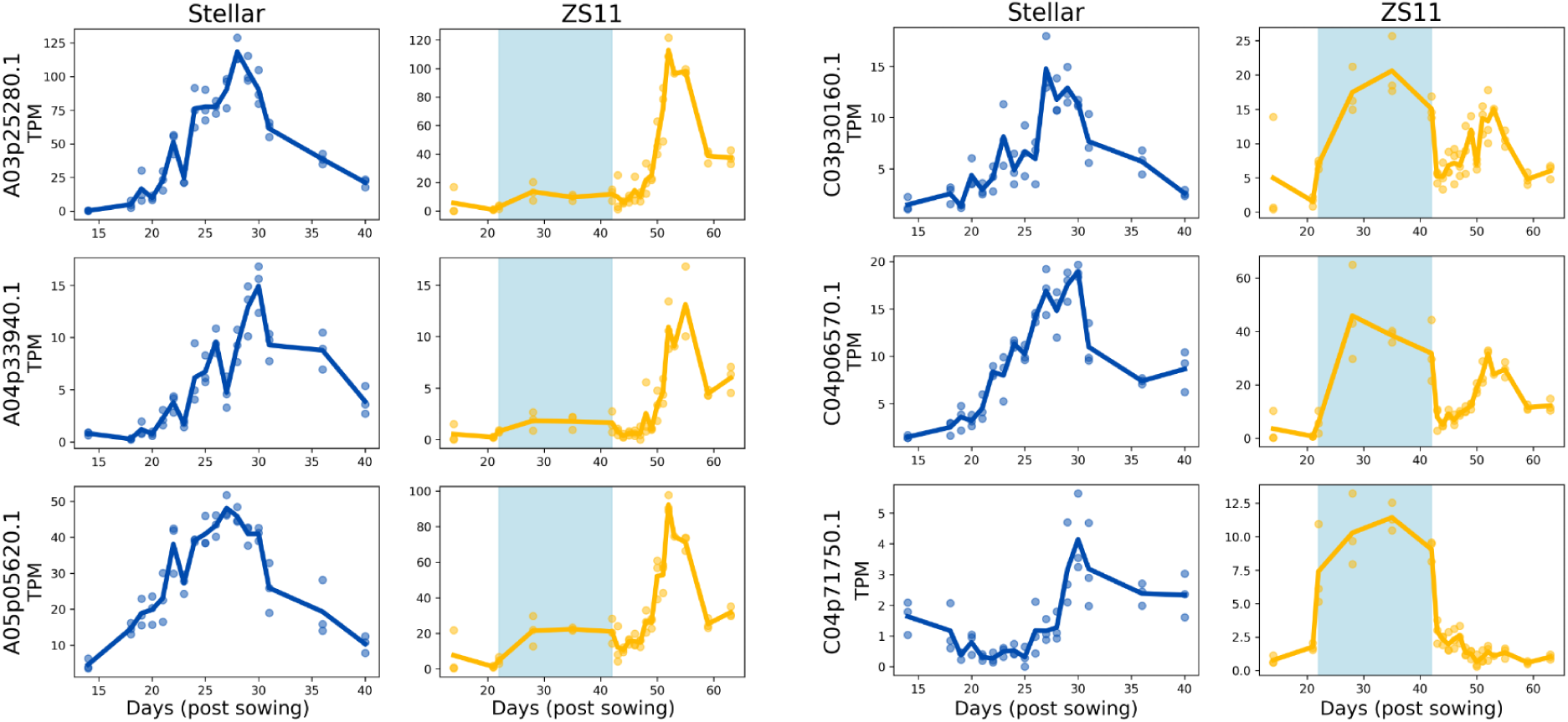
*SOC1* orthologues in ZS11 have differences in expression profiles under short-day, cold treatment. The plots show the expression of *SOC1* orthologues in both Stellar (dark blue) and ZS11 (yellow) cultivars, with the short-day, cold period highlighted in blue in the ZS11 plots. The details for these timeseries are shown in Figure 1. Briefly, Stellar undergoes the floral transition at day 30 while ZS11 transitions at day 55. All *SOC1* orthologues follow similar dynamics in Stellar, with an increase until the apex becomes floral and a decrease afterwards, albeit at different expression levels. However, in ZS11, orthologues have varying degrees of upregulation during short-day, cold treatment, indicating regulatory divergence. The three C genome orthologues have their expression maxima during that period, which does not coincide with the floral transition.

The *SOC1* paralogues also exhibit variation in expression level; this is consistent between the two cultivars. *Bna.SOC1.A03* and *Bna.SOC1.A05* have the highest expression, while *Bna.SOC1.C04b* and *Bna.SOC1.A04* have the lowest.

Overall, the ZS11 timeseries shows that orthologues of *SOC1* in *B. napus* have differences in their dynamics and therefore likely their regulation. It is, however, unclear if this is due to short-day, cold treatment or due to differences between the two cultivars.

### 3.4 *SOC1* paralogues have diverged in their regulation under cold temperature conditions in *B. napus*

The analysis of inferred networks in *B. napus* cv. ZS11 suggests differences in regulation of the six orthologues of the floral integrator, *SOC1*. This is evident from the differences in their expression dynamics following the start of short-day, cold treatment. However, their expression dynamics and inferred regulators appear similar in *B. napus* cv. Stellar. To investigate whether these observations are the result of differences between cultivars or in response to environmental change, we grew *B. napus* cv Stellar plants under the same short-day, cold conditions. The sampled timepoints are detailed in Figure 6 (a). We observed a delayed floral transition under short-day, cold treatment compared to normal conditions with the floral shoot apical meristem visible on day 37 (Supplemental Figure S6), compared to day 30 in the earlier timeseries (Figure 1).

**Figure 6:**
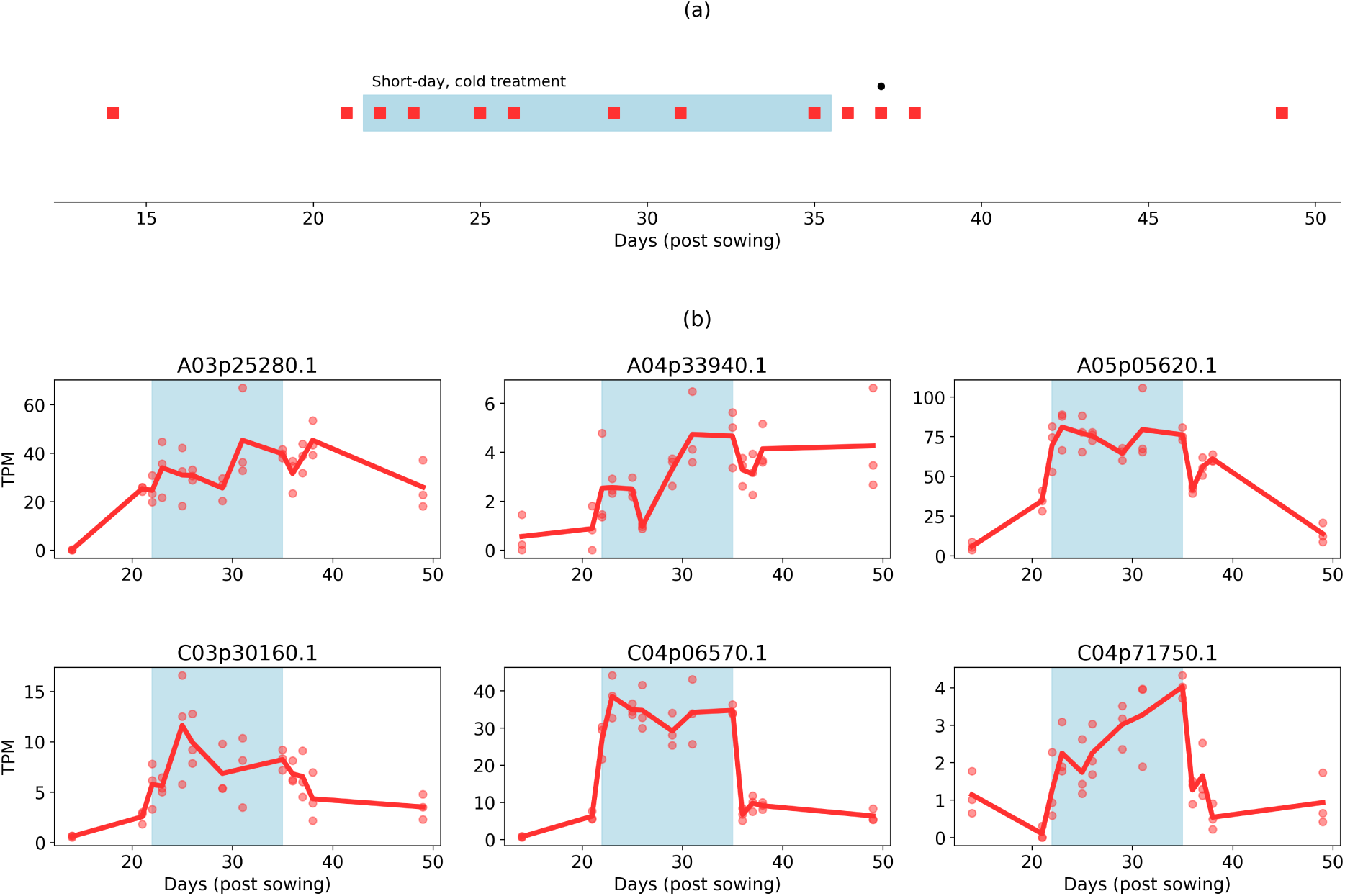
*SOC1* orthologues exhibit divergent expression dynamics in response to short-day, cold conditions in Stellar. In order to ascertain if the divergence expression dynamics in *SOC1* orthologues was due to the environmental change or differences between cultivars, we grew Stellar plants with the same short-day, cold treatment as ZS11 plants. (a) The red markers show when SAMs were sampled for RNA-seq. Plants given two weeks of short-day, cold treatment transitioned at day 37 instead of day 30, observed in previous timeseries without the treatment (Figure 1). Supplementary Table 5 contains detailed alignment statistics for the timeseries. (b) *SOC1* orthologues shows divergent expression dynamics, with some orthologues exhibiting upregulation when plants are moved to short-day, cold conditions. By comparing the expression dynamics under normal conditions in Stellar (Figure 5), we infer that the expression divergence in *SOC1* orthologues is not due to differences between cultivars but due to differences in regulation.

Figure 6 (b) shows expression levels of the *SOC1* orthologues in this new timeseries. Compared with the expression data from the earlier Stellar timeseries (Figure 5), *SOC1* orthologues have clear differences in their dynamics, indicating that the inferred differences in *SOC1* regulation are indeed in response to short-day, cold treatment rather than being cultivar specific. *Bna.SOC1.A03* was the highest expressed orthologue when both Stellar and ZS11 underwent the floral transition (as shown in Figure 5), however, *Bna.SOC1.A05* becomes the highest expressed orthologue when plants are exposed to short day, cold conditions. When plants are returned to warm, long day conditions expression levels return to *Bna.SOC1.A03* being higher than *Bna.SOC1.A05* by day 49. The expression of *Bna.SOC1.A04* is low in all sampled datasets. The three C genome orthologues *Bna.SOC1.C03*, *Bna.SOC1.C04a* and *Bna.SOC1.C04b* have their expression maxima during the short-day, cold treatment. These observations are similar to the dynamics observed in the ZS11 timeseries, (Figure 5) and suggest that orthologues of *SOC1* have diverged in their regulation, and that this, potentially subfunctionalised, response is only exhibited under short-day, cold conditions.

The genes that are upregulated under short-day, cold conditions could be performing different biochemical functions from Arabidopsis *SOC1*. However, multiple sequence alignment of protein sequences shows high sequence conservation (Supplemental Figure S7 and Supplemental Table 6). The functional MADS domain, constituting the first 57 amino acids from the N-terminal that is required for translocation of *SOC1* to the nucleus [37] is identical to the Arabidopsis *SOC1* in all *B. napus* orthologues, except in *Bna.SOC1.A05* and *Bna.SOC1.C04a*, which have a substitution from Glycine (G) to Alanine (A) at position 52. Therefore, whilst we identified changes in regulation that are suggestive of subfunctionalisation, we did not find evidence of functional divergence among the *B. napus SOC1* orthologues.

To determine if any known regulators could be causing changes in expression, we investigated the effect of short-day, cold treatment on upstream regulators of *SOC1* that are known to directly bind to the *SOC1* promoter before the floral transition. These proteins (Supplementary Table 7) have been narrowed down from a wider list of putative and known *SOC1* regulators to exclude indirect regulators, regulators without a known binding site and post-floral transition regulators obtained from a previous study [68] and the FLOR-ID database [9]. Orthologues of one of these regulators, *CIRCADIAN CLOCK ASSOCIATED 1* (*CCA1*) are, interestingly, only expressed under short-day, cold conditions in *B. napus* (Figure 7 (a)).

**Figure 7:**
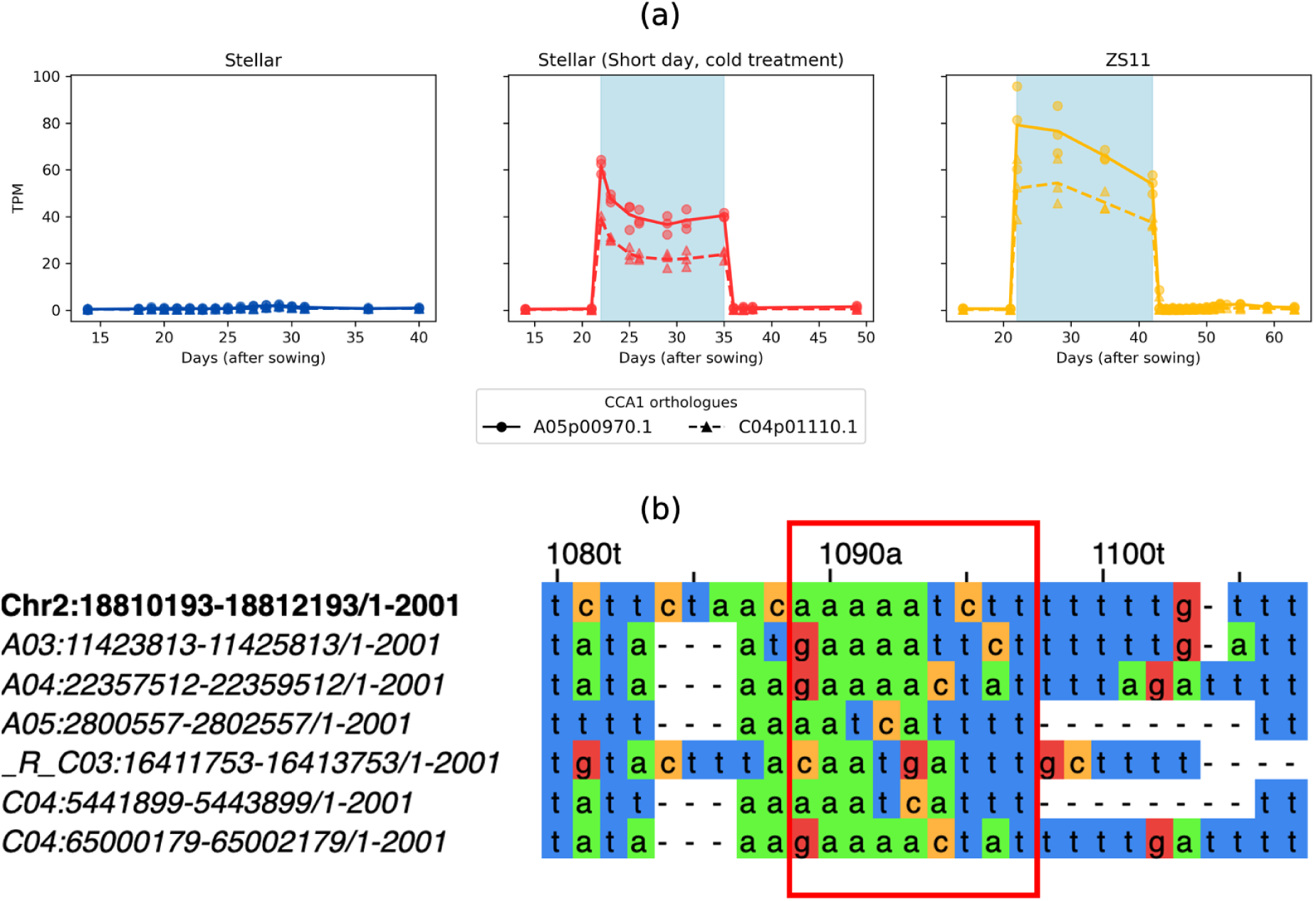
Orthologues of *CCA1* are upregulated under short-day, cold conditions. *CIRCADIAN CLOCK ASSOCIATED* (*CCA1*) is an upstream regulator of *SOC1* in Arabidopsis [43]. (a) The expression data shows that orthologues of *CCA1* are significantly upregulated in both cultivars of *B. napus*, when plants are subjected to short-day, cold conditions. (b) Multiple sequence alignment of upstream regions of Arabidopsis *SOC1* and its six orthologues in *B. napus* shows variation in the binding sequence for *CCA1* in the promoter regions. The labels show the exact coordinates of the promoter regions used for alignment from the TAIR 10 reference (Chr 2; for Arabidopsis) [36] and Darmor v10 [60] (A03, A04, A05, C03 and C04).

There are two orthologues of *CCA1* in *B. napus* with similar expression dynamics and levels. The expression of these orthologues is very low in both Stellar and ZS11 under normal conditions, however, they are rapidly upregulated as soon as plants are shifted to short-day, cold conditions. *SOC1* expression has been shown to be significantly repressed in a *CCA1* overexpressor line and ChIP analysis has shown that *CCA1* transcription factor binds to a site upstream of *SOC1* [43]. Furthermore, a study in related *Brassicaceae* has indicated that there is high variation in the binding sites for known regulators among *SOC1* orthologues [68].

Based on these reports, we isolated 2000 bp upstream promoter regions and performed multiple sequence alignment analysis. Focussing on the promoter region previously determined to be the binding site for *CCA1* within Arabidopsis, our analysis shows variation among all *SOC1* orthologues for this site, indicating that binding for *CCA1* could be disrupted. Sequences have variation in the trailing ‘TCTT’ sequence (Figure 7 (b)), with *Bna.SOC1.A03* having the closest match to the Arabidopsis site.

While the transcription factor binding site disruption on its own is not strong evidence, together with the expression pattern of *CCA1* orthologues, we speculate that *CCA1* upregulation might be responsible for the differences in expression of orthologues of *SOC1* in *B. napus*. The *Bna.SOC1.A03* promoter has the binding site most similar to Arabidopsis and is kept repressed under short-day, cold conditions, while all other *SOC1* orthologues have varying degrees of upregulation coupled with variations in the *CCA1* binding site.

Our data were collected under both changing photoperiod and temperature conditions.As *CCA1* is controlled by the circadian clock, either of these environmental changes, or their combination, could be the reason for its upregulation. *CCA1* has been reported to undergo alternative splicing under low temperatures and has been posited to be a link between circadian pathway and cold response [53], we therefore hypothesised that changes in expression dynamics may be independent of photoperiod change.

To determine the stimulus that leads to the changes in dynamics of *CCA1* and *SOC1* orthologues, we generated two further shoot apical meristem RNA-seq timeseries from *B. napus* cv. Stellar plants. We exposed one set of plants to two weeks of cold temperature treatment while keeping the photoperiod constant, while the other set received constant temperature treatment under short-day conditions. The sampled timepoints, along with the results are shown in Figure 8. Changes in dynamics of *SOC1* orthologues are present under cold temperature, even when photoperiod was kept constant. *CCA1* orthologues are also upregulated in the same set of plants, illustrating the effect of cold on gene expression. The plants that received a short-day treatment show no such change, confirming that cold temperature was responsible for upregulation of both *SOC1* and *CCA1* orthologues in *B. napus*.

**Figure 8:**
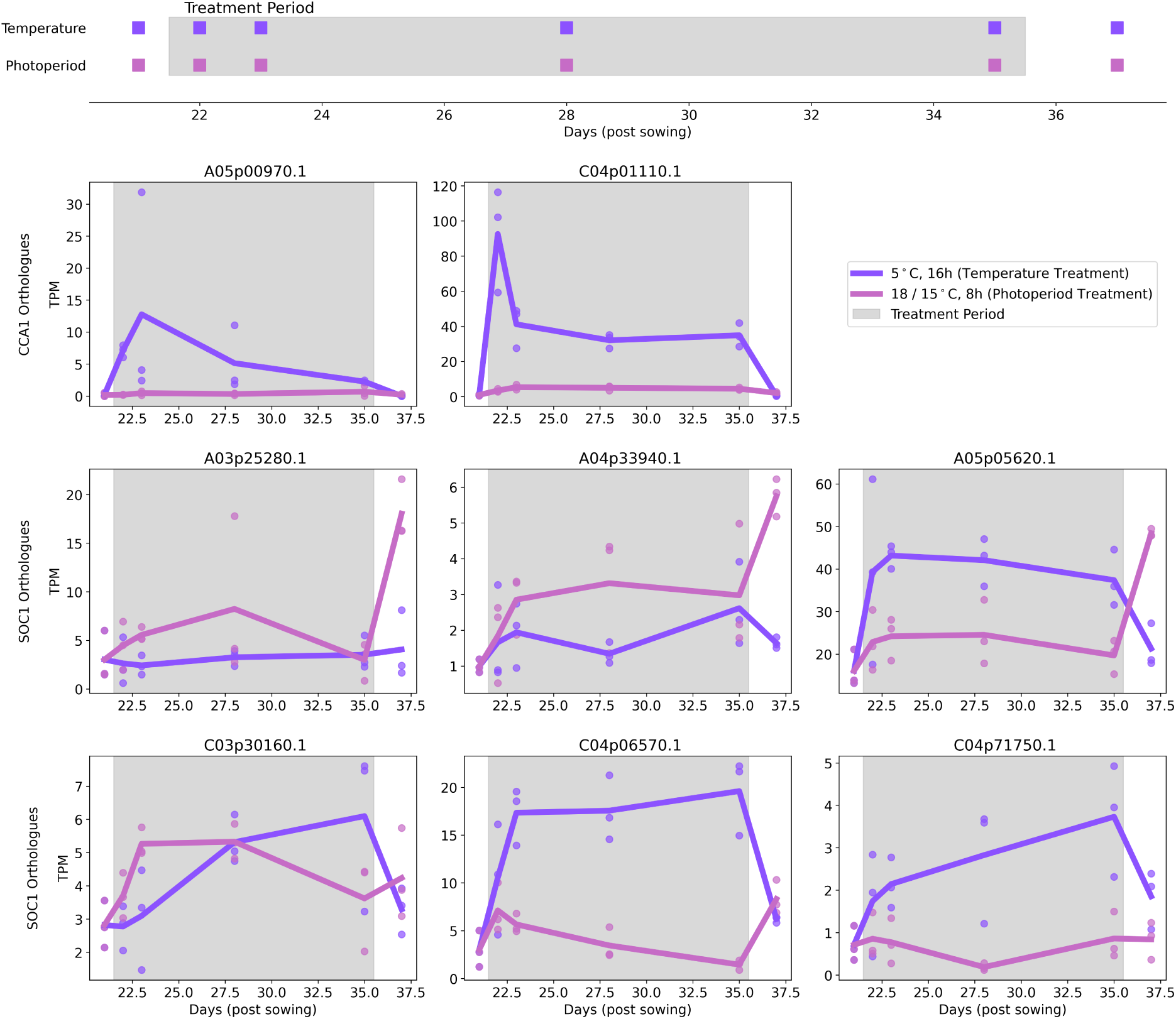
*SOC1* orthologues show divergence in their expression dynamics and *CCA1* orthologues are upregulated in response to change in temperature. We subjected *B. napus* cv. Stellar plants to either a change in temperature or photoperiod and sampled SAMs to observe expression dynamics of orthologues of *SOC1* and its upstream regulator in Arabidopsis, *CCA1*. We observe that during treatment period, the expression of *CCA1* orthologues is higher in plants that were subjected to cold conditions compared to plants that were subjected to short-day treatment. Similarly, *SOC1* orthologues *Bna.SOC1.A05*, *Bna.SOC1.C04a* and *Bna.SOC1.C04b* exhibit higher expression in plants that were given cold treatment. *Bna.A03.SOC1*, which is the highest expressed copy under normal conditions is repressed under cold conditions. This confirms that the regulatory divergence observed among *SOC1* orthologues is in response to cold temperature conditions.

## 4 Discussion

Research into the control of flowering time in Arabidopsis has provided a detailed understanding of the GRN controlling the timing of this transition. There is significant interest in translating this knowledge into crop species to aid the development of varieties to ensure food security in the face of rapid climate change. Due to its close evolutionary relationship with Arabidopsis, *B. napus* is a prime candidate for such translational research. We gathered detailed data on expression dynamics of genes throughout plant development and used computational techniques to compare expression dynamics of orthologues between Arabidopsis and *B. napus* and infer regulatory relationships among flowering time orthologues in *B. napus*.

Using curve registration, we show that orthologues of Arabidopsis genes in polyploid *B. napus* retain similar expression dynamics, and likely similar regulation on a whole genome level. The GRN controlling flowering time also has similarities to Arabidopsis, however, preferential retention of flowering time genes allows individual paralogues to diverge in expression, while still maintaining the required structure of the regulatory framework. This allows plasticity in the system, aiding in adaptations and giving rise to the wide diversity of cultivar types. Previous research has shown that flowering time genes are preferentially retained and expressed [27] [62] and individual paralogues of *FLC* have been shown to have divergence in expression dynamics [12]. Our results build on these works and statistically show that it is a more widespread phenomenon among orthologues of flowering time genes, lending further support to the idea that each individual orthologue needs to be studied for functional divergence to select useful breeding targets [61].

We discover and present one such example of probable subfunctionalisation in detail. We did not find any indication of functional divergence between the six orthologues of the key floral integrator gene, *SOC1*, in *B. napus* based on protein sequence analysis. The orthologues have different expression levels but similar expression dynamics under long-day, normal temperature conditions, with all genes having their expression maxima coinciding with the floral transition. Under cold temperature, however, the expression dynamics change, with rapid upregulation of some paralogues. Assuming all *SOC1* orthologues are still performing the same function, this likely means that some orthologues are responsible for floral transition under cold temperature, while others are more important under warmer conditions. Similar conclusions can be drawn from the data from a long-day vernalisation experiment conducted by Matar *et al*. in winter-type *B. napus* cultivars, where the expression of only two orthologues of *SOC1*, the paralogue on chromosome A05 and its homologue on C04 in the Express617 genome, allowed the plants to floral transition under cold conditions [48]. *CCA1* upregulation under cold, and likely disruption in its ability to repress some *SOC1* orthologues due to variation observed in binding site sequences provides some explanation for the divergence observed. Previous studies have also reported differences among promoter regions of *SOC1* orthologues [47] [68], indicating that there are likely more regulatory divergences warranting further investigations.

## Author Contributions

GSS did the sampling, planned, designed and performed all the data analysis. SB and RW provided sampling support. HW provided bioinformatics support. RW and RJM conceived and supervised the project. GSS prepared the figures and wrote the first draft of the manuscript. All authors contributed to the writing and editing of the final version manuscript.

## Data and code availability

The RNA-seq data from this study has been deposited in the European Nucleotide Archive (ENA) under projects PRJEB106353, PRJEB106354 and PRJEB106824. The custom python script used to perform reciprocal blast search is available on zenodo: https://doi.org/10.5281/zenodo.17631422. Full set of curve registration results are available in Supplemental File 1. The input expression data and outputs for all inferred networks are available in Supplemental File 2.

## Acknowledgements

We are grateful to Thomas Lock, Judith Irwin, Lorelei Bilham, Steve Penfield and Carmel O’Neill for their generous help with sampling some of the timepoints in the timeseries presented in this study. We also acknowledge support by the NBI Research Computing and Horticultural Services.

## Supplementary Information

### Supplementary Figures

**Figure S1:**
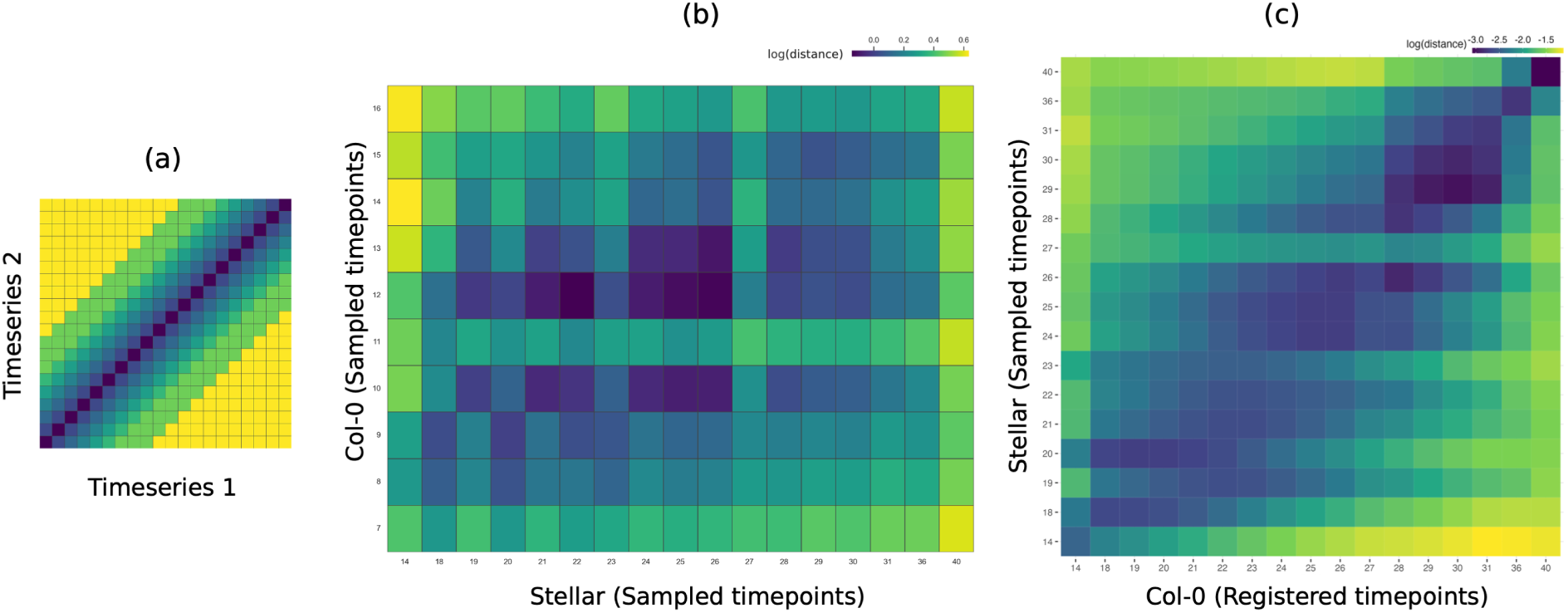
Curve registration facilitates comparison of RNA-seq time courses across timescales. The heatmaps show pairwise distance between gene expression at all timepoints between the *B. napus* cv. Stellar and Arabidopsis Col-0 timeseries for flowering time genes. (a) A comparison of pairwise distances between two timeseries with similar progression will result in a plot with minimum distances along the diagonal, represented by dark blue colours while timepoints further away would be maximally distant, shown by green-yellow colours. (b) Pairwise distances with just normalised expression data (scaled to be between 0 and 1) between Stellar and Col-0 timeseries. This comparison does not show any similarity in progression between the time courses. (c) Using greatR [34] to register the Col-0 timeseries against the reference Stellar, reveals a diagonal, showing that there are similarities between the two timeseries.

**Figure S2:**
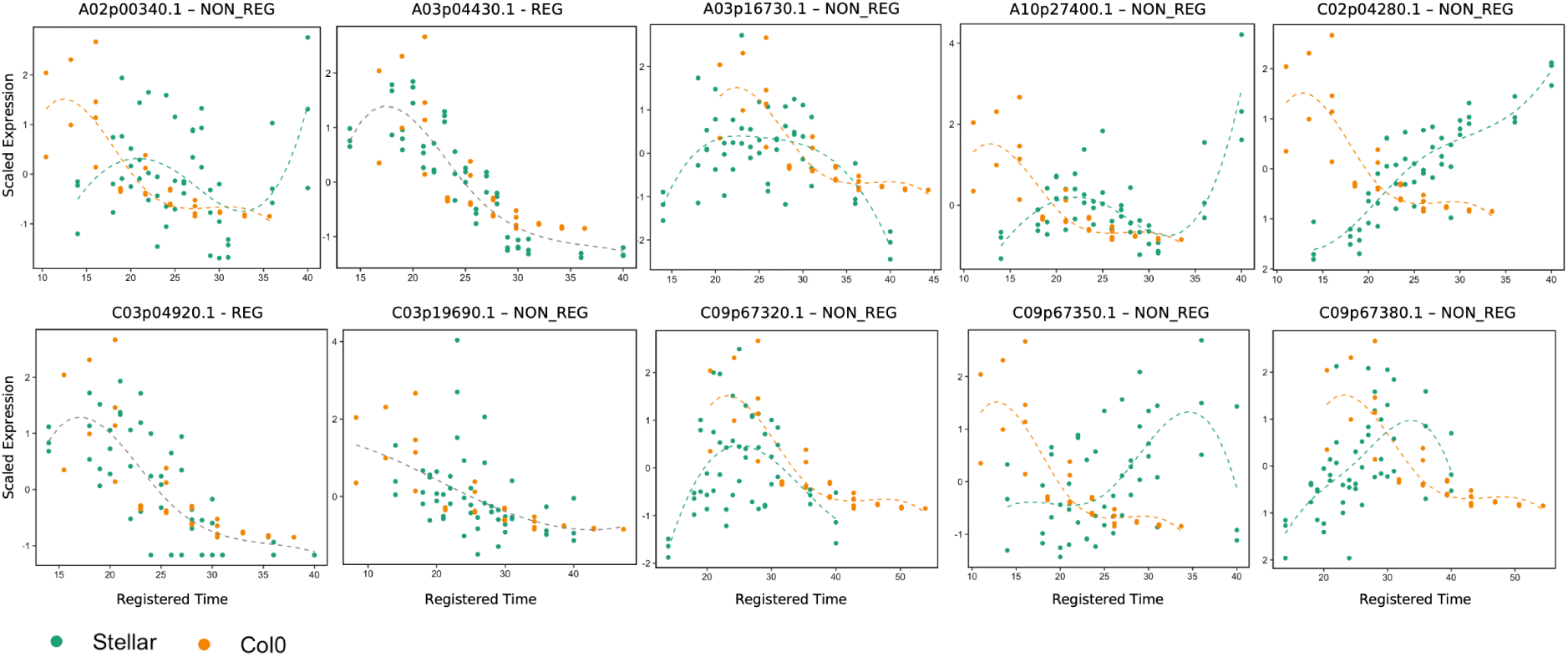
Curve registration highlights the divergence in expression dynamics among *FLC* orthologues in *B. napus* cv. Stellar. *B. napus* Darmor v10 reference genome [60] contains 10 orthologues for the Arabidopsis *FLOWERING LOCUS C* (*FLC*). Only 3 orthologues A03p04430.1, C03p04920.1 and C03p19690.1 show dynamics similar to the Arabidopsis *FLC* and hence are ‘registered’. The plot titles show the differences in calculated Bayesian Information Criterion (BIC) values, and the corresponding stretch and shift factors applied to that gene by greatR [34]. The ‘non-registered’ orthologues have clearly different dynamics to the Arabidopsis *FLC*. For instance, C02p04280.1 and C09p67380.1 increase in expression as the plant approaches floral transition, which is exactly opposite to the dynamics of the Arabidopsis *FLC* gene.

**Figure S3:**
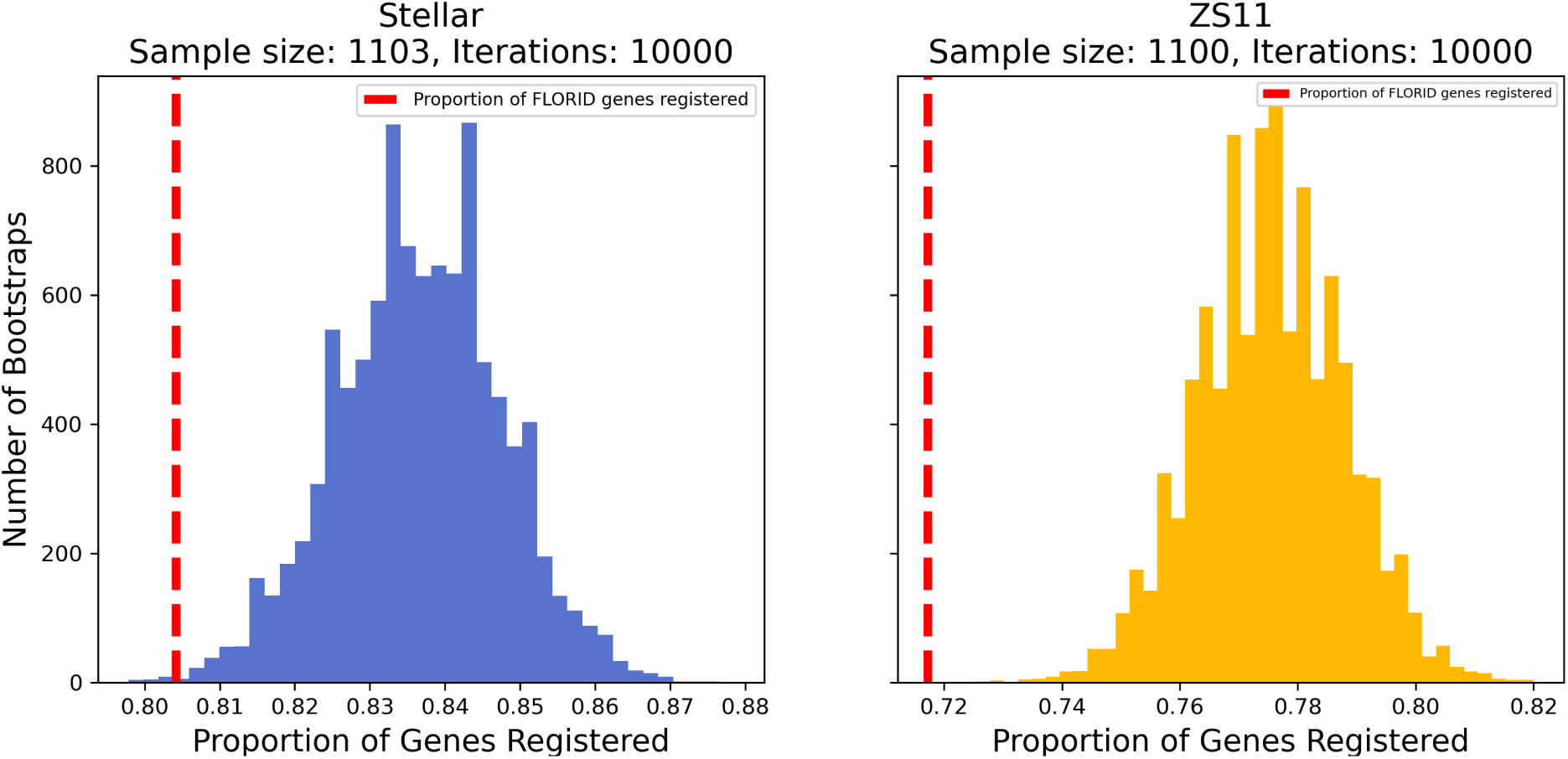
The lower proportion of registration observed among orthologues of flowering time genes is unlikely to be by a random chance. The plots show results from a bootstrapping experiment in which samples of genes of sizes 1103 and 1100 for Stellar and ZS11 timeseries were sampled with replacement from a set of all genes for 10,000 iterations. The sample sizes are equal to the number of flowering genes that were expressed in both cultivars, based on filtering criteria of standard deviation *>* 0, used by greatR [34]. The red bar shows the proportion of flowering time orthologues registered, represented on the x-axis. The plots show that the red line falls outside the distributions obtained following sampling, showing that it is unlikely that the lower proportion of registration to Arabidopsis orthologues observed for the set of flowering time genes is by random chance.

**Figure S4:**
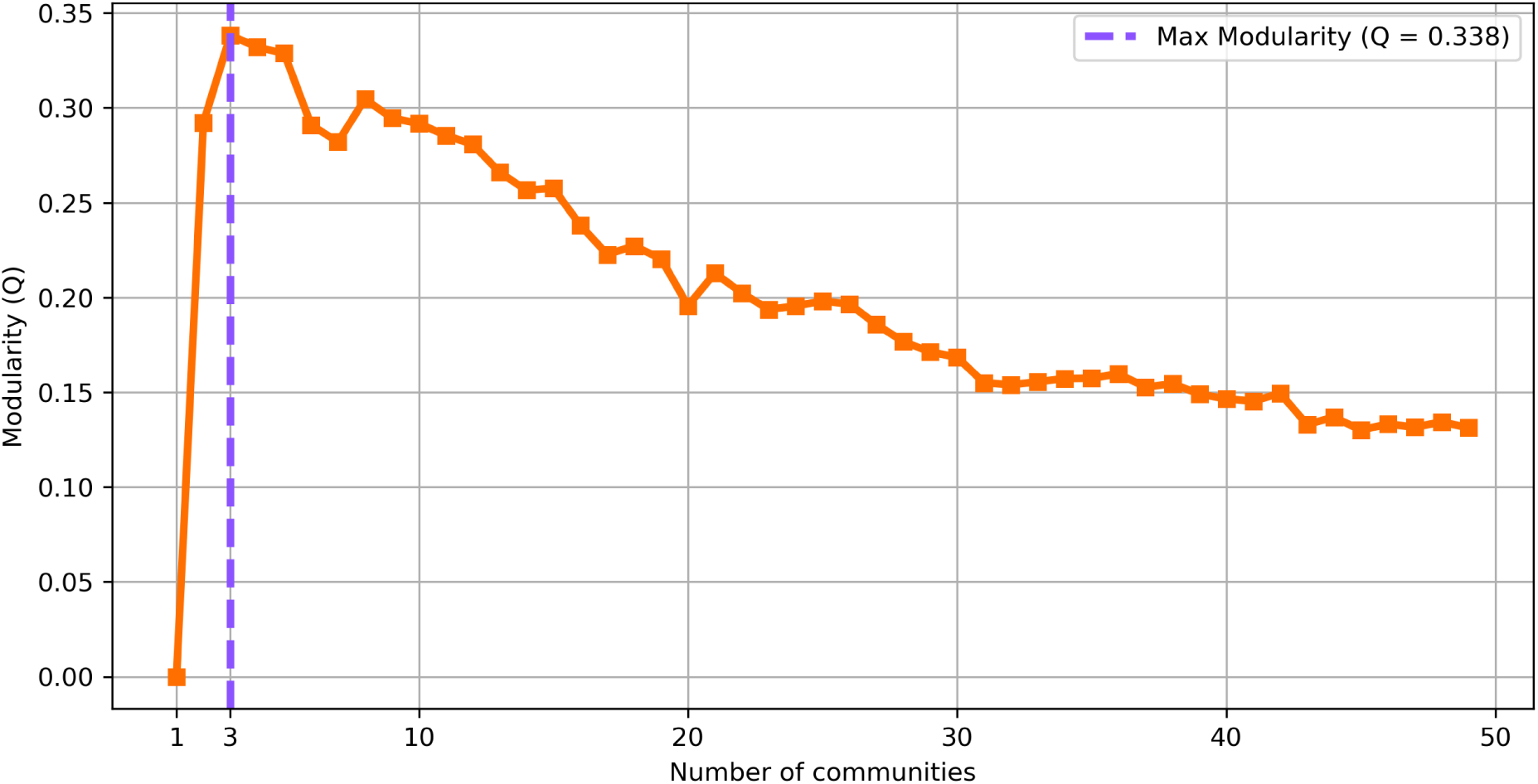
ZS11 network can be divided into three communities using the maximum modularity method. The plot shows the calculation of the modularity metric for dividing the network inferred from the ZS11 timeseries into various number of communities. The maximum modularity occurs when number of communities equals 3, so it was selected as the optimum number to divide the network.

**Figure S5:**
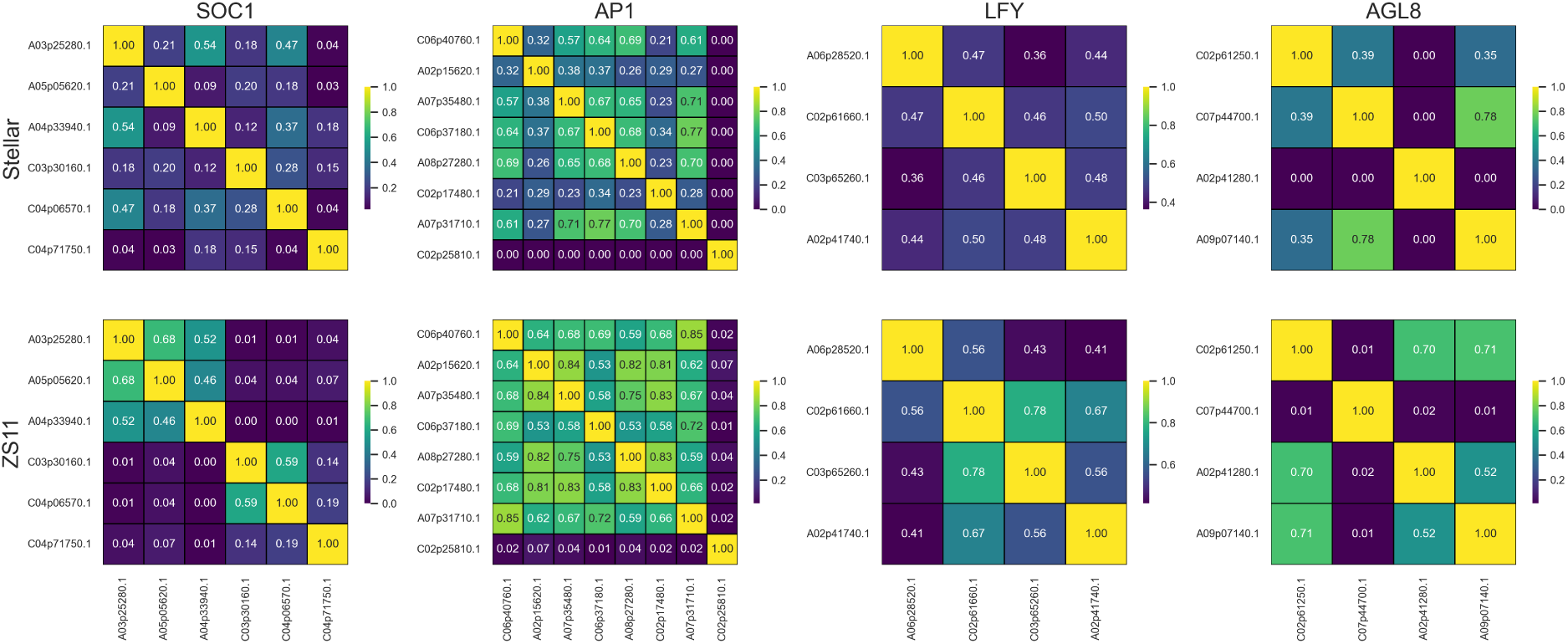
Orthologues of *SOC1* are unique among floral integrators to have major differences in upstream regulators among different paralogues between the two *B. napus* networks of flowering time genes. The heatmaps show pairwise Rank-Biased overlap [77] scores for regulators of orthologues of floral integrators *SOC1*, *AP1*, *LFY* and *AGL8*. Each cell in the heatmap represents the overlap between the ranking of upstream regulators of the two genes, with diagonals, representing the comparison of a gene with itself, having the maximum value of 1.0. The first row corresponds to data from the network inferred from Stellar data, while second row is for ZS11. The heatmaps show that unlike for *AP1*, *LFY* and *AGL8*, orthologues of *SOC1* appear to from groups with clear differences in regulation in the network inferred from ZS11 timeseries. These differences are not as stark in the network inferred from the Stellar timeseries data.

**Figure S6:**
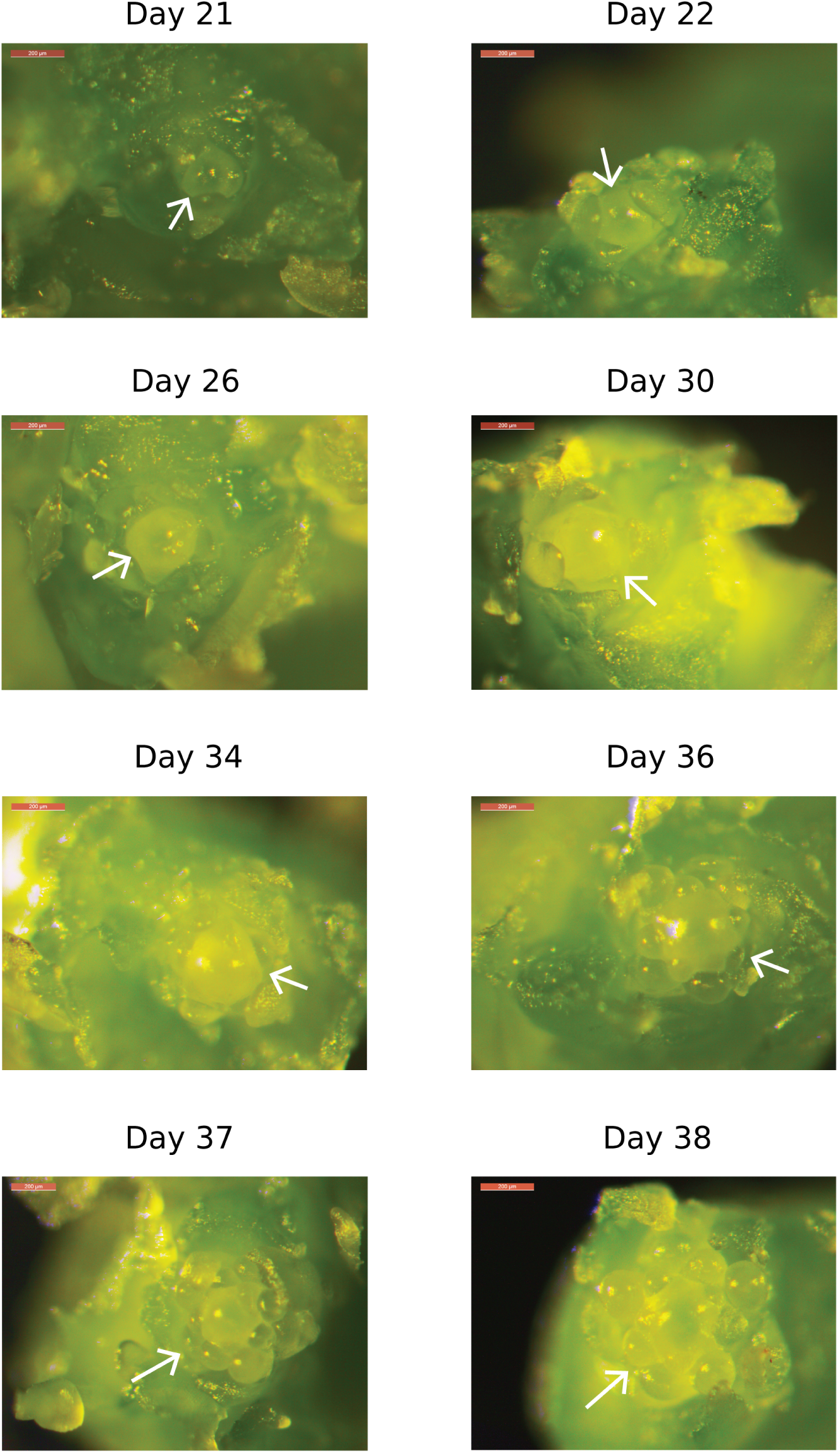
The Stellar shoot apical meristem (SAM) turns floral at day 37 when plants were given two weeks of short-day, cold treatment (5 *^◦^*C, 8-hour photoperiod). Images of the dissected SAMs under a light microscope at various time-points. The SAM, indicated by the white arrow, appears clearly vegetative until day 34. Some floral primordia begin appearing on day 36, and day 37 is when floral primordia become clearly visible, marking the floral transition.

**Figure S7:**
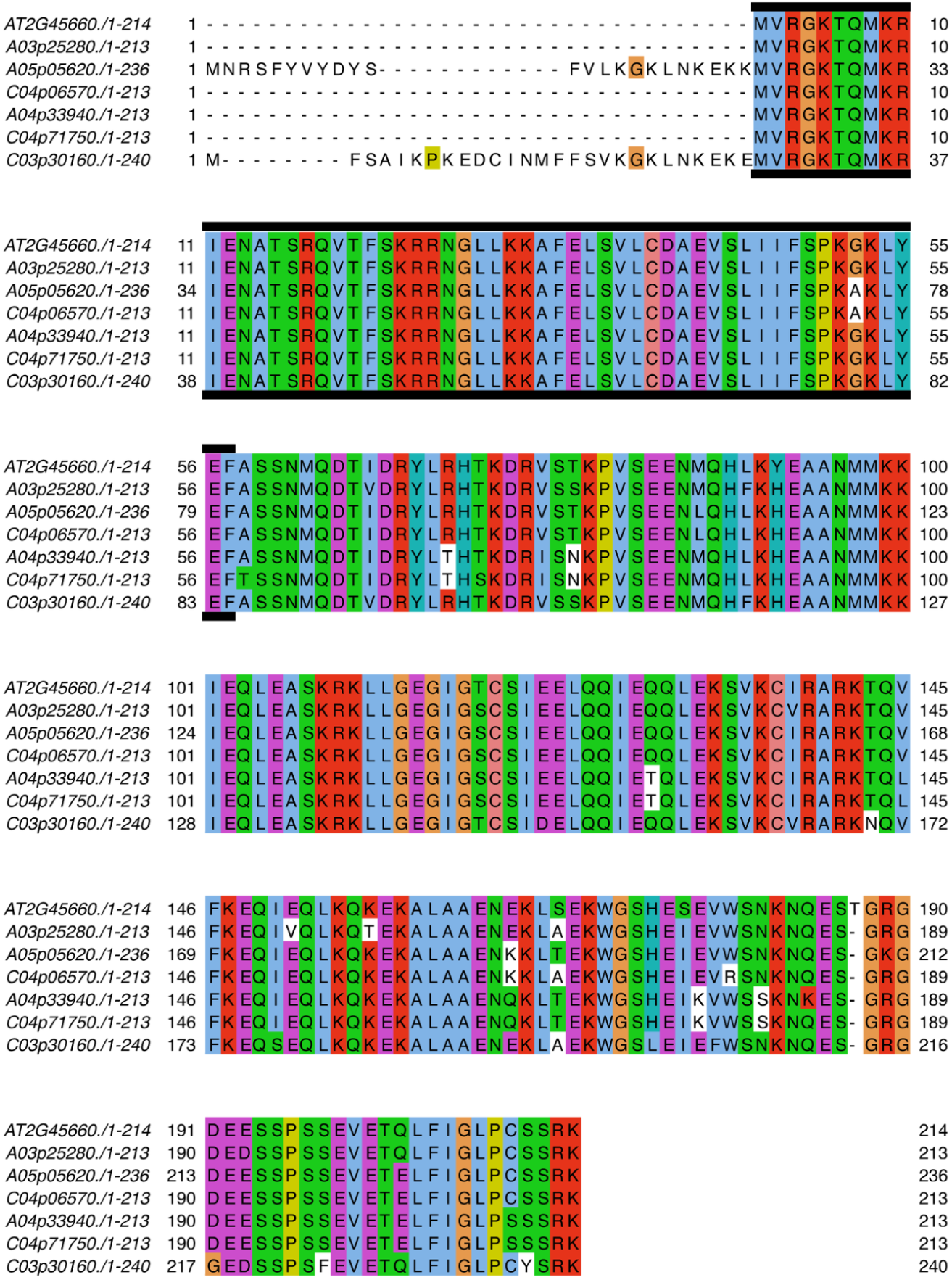
Protein sequences for Arabidopsis *SOC1* and its *B. napus* orthologues are highly conserved. Multiple sequence alignment shows that all *SOC1* proteins in *B. napus* have highly conserved sequences. The MADS domain, highlighted in the figure, is also highly conserved, except in two paralogues with one substitution from Glycine (G) to Alanine (A) at position 52. Alanine has a bulkier methyl group compared to Glycine, however, structural information is needed to determine any functional effects.

### Supplementary Tables

**Table 1:** Alignment statistics for *B. napus* cv. Stellar RNA-seq timeseries. Samples in Stellar timeseries, as shown in Figure 1 show consistent alignment rate to the Darmor v10 reference genome [60], with an average of 89.99 %. In every sample, majority of reads are uniquely mapped, with an average of 73.60 %.

| Timepoint<br>(Days) | Replicate | Total<br>Reads<br>(Raw) | Total<br>Reads<br>(QC<br>passed) | Mapped<br>Reads<br>(Total) | Mapped<br>Reads<br>(Unique) | Mapped<br>Reads<br>(Multi) | Alignment<br>Rate<br>(Total,<br>%) | Alignment<br>Rate<br>(Unique,<br>%) |
| --- | --- | --- | --- | --- | --- | --- | --- | --- |
| 14 | 1 | 29466748 | 28758229 | 26250311 | 21450623 | 4799688 | 91.28 | 74.59 |
| 14 | 2 | 24666990 | 23939901 | 22244659 | 18205509 | 4039150 | 92.92 | 76.05 |
| 14 | 3 | 24219439 | 23540721 | 20695184 | 16720576 | 3974608 | 87.91 | 71.03 |
| 18 | 1 | 27746132 | 26994954 | 24039328 | 19672135 | 4367193 | 89.05 | 72.87 |
| 18 | 2 | 33326240 | 32381227 | 30014138 | 24840379 | 5173759 | 92.69 | 76.71 |
| 18 | 3 | 27514526 | 26844350 | 23813388 | 19441293 | 4372095 | 88.71 | 72.42 |
| 19 | 1 | 30157849 | 29446093 | 25690963 | 20873799 | 4817164 | 87.25 | 70.89 |
| 19 | 2 | 27995140 | 27481926 | 25596698 | 21166876 | 4429822 | 93.14 | 77.02 |
| 19 | 3 | 25437387 | 24794130 | 22219958 | 18092271 | 4127687 | 89.62 | 72.97 |
| 20 | 1 | 30370848 | 29405460 | 26043924 | 21347396 | 4696528 | 88.57 | 72.60 |
| 20 | 2 | 30099757 | 29388288 | 27204557 | 22345602 | 4858955 | 92.57 | 76.04 |
| 20 | 3 | 27622931 | 26912498 | 24237255 | 19923191 | 4314064 | 90.06 | 74.03 |
| 21 | 1 | 30529239 | 29805535 | 26592738 | 21588327 | 5004411 | 89.22 | 72.43 |
| 21 | 2 | 29794316 | 29136562 | 26831615 | 21922714 | 4908901 | 92.09 | 75.24 |
| 21 | 3 | 36161871 | 35106563 | 30734332 | 25166413 | 5567919 | 87.55 | 71.69 |
| 22 | 1 | 32216530 | 31394820 | 27383769 | 21996099 | 5387670 | 87.22 | 70.06 |
| 22 | 2 | 27599676 | 26998808 | 24780379 | 20283358 | 4497021 | 91.78 | 75.13 |
| 22 | 3 | 30980902 | 29687457 | 25480607 | 21084379 | 4396228 | 85.83 | 71.02 |
| 23 | 1 | 26562465 | 25762479 | 23996188 | 19771662 | 4224526 | 93.14 | 76.75 |
| 23 | 2 | 28097869 | 27432127 | 25438002 | 20846655 | 4591347 | 92.73 | 75.99 |
| 23 | 3 | 29856661 | 28439215 | 24767469 | 20450559 | 4316910 | 87.09 | 71.91 |
| 24 | 1 | 24302179 | 23720233 | 22227620 | 18323458 | 3904162 | 93.71 | 77.25 |
| 24 | 2 | 25725616 | 25002390 | 22554092 | 18347106 | 4206986 | 90.21 | 73.38 |
| 24 | 3 | 34036988 | 33161191 | 29517525 | 24103629 | 5413896 | 89.01 | 72.69 |
| 25 | 1 | 25441290 | 24739891 | 22925859 | 18769779 | 4156080 | 92.67 | 75.87 |
| 25 | 2 | 25364313 | 24703104 | 22616725 | 18484703 | 4132022 | 91.55 | 74.83 |
| 25 | 3 | 34674556 | 33787709 | 28811580 | 23528903 | 5282677 | 85.27 | 69.64 |
| 26 | 1 | 27883642 | 27133476 | 24622563 | 20091788 | 4530775 | 90.75 | 74.05 |
| 26 | 2 | 26767602 | 26016735 | 24070235 | 19627849 | 4442386 | 92.52 | 75.44 |
| 26 | 3 | 30754646 | 29893009 | 26058349 | 21372331 | 4686018 | 87.17 | 71.50 |
| 27 | 1 | 29092425 | 28297242 | 25591451 | 20748923 | 4842528 | 90.44 | 73.32 |
| 27 | 2 | 27636798 | 26948440 | 24168101 | 19537868 | 4630233 | 89.68 | 72.50 |
| 27 | 3 | 27314762 | 26627251 | 23070553 | 18893948 | 4176605 | 86.64 | 70.96 |
| 28 | 1 | 30368320 | 29648974 | 27310613 | 22373761 | 4936852 | 92.11 | 75.46 |
| 28 | 2 | 25388514 | 24712203 | 22165757 | 18035580 | 4130177 | 89.70 | 72.98 |
| 28 | 3 | 35986575 | 34972251 | 29791105 | 24352870 | 5438235 | 85.18 | 69.63 |
| 29 | 1 | 29019928 | 28365045 | 26366028 | 21633502 | 4732526 | 92.95 | 76.27 |
| 29 | 2 | 26945703 | 26330055 | 23975406 | 19563169 | 4412237 | 91.06 | 74.30 |
| 29 | 3 | 32549747 | 31559450 | 26980658 | 22142982 | 4837676 | 85.49 | 70.16 |
| 30 | 1 | 31955891 | 31242074 | 28555816 | 23213624 | 5342192 | 91.40 | 74.30 |
| 30 | 2 | 28568289 | 27908034 | 25435294 | 20680682 | 4754612 | 91.14 | 74.10 |
| 30 | 3 | 35836730 | 34892582 | 29785513 | 24392437 | 5393076 | 85.36 | 69.91 |
| 31 | 1 | 32110567 | 31256305 | 29136417 | 23968576 | 5167841 | 93.22 | 76.68 |
| 31 | 2 | 25037174 | 24462520 | 21993226 | 17886998 | 4106228 | 89.91 | 73.12 |
| 31 | 3 | 36175210 | 35194846 | 29940097 | 24560176 | 5379921 | 85.07 | 69.78 |
| 36 | 1 | 29711573 | 28943845 | 26905093 | 22156305 | 4748788 | 92.96 | 76.55 |
| 36 | 2 | 24666008 | 24092183 | 21873027 | 17825171 | 4047856 | 90.79 | 73.99 |
| 36 | 3 | 36570825 | 35493107 | 31326379 | 25658076 | 5668303 | 88.26 | 72.29 |
| 40 | 1 | 24759987 | 24082704 | 22577507 | 18681864 | 3895643 | 93.75 | 77.57 |
| 40 | 2 | 29149451 | 28423345 | 26149052 | 21360606 | 4788446 | 92.00 | 75.15 |
| 40 | 3 | 38484447 | 37477496 | 33408680 | 27256602 | 6152078 | 89.14 | 72.73 |

**Table 2:**
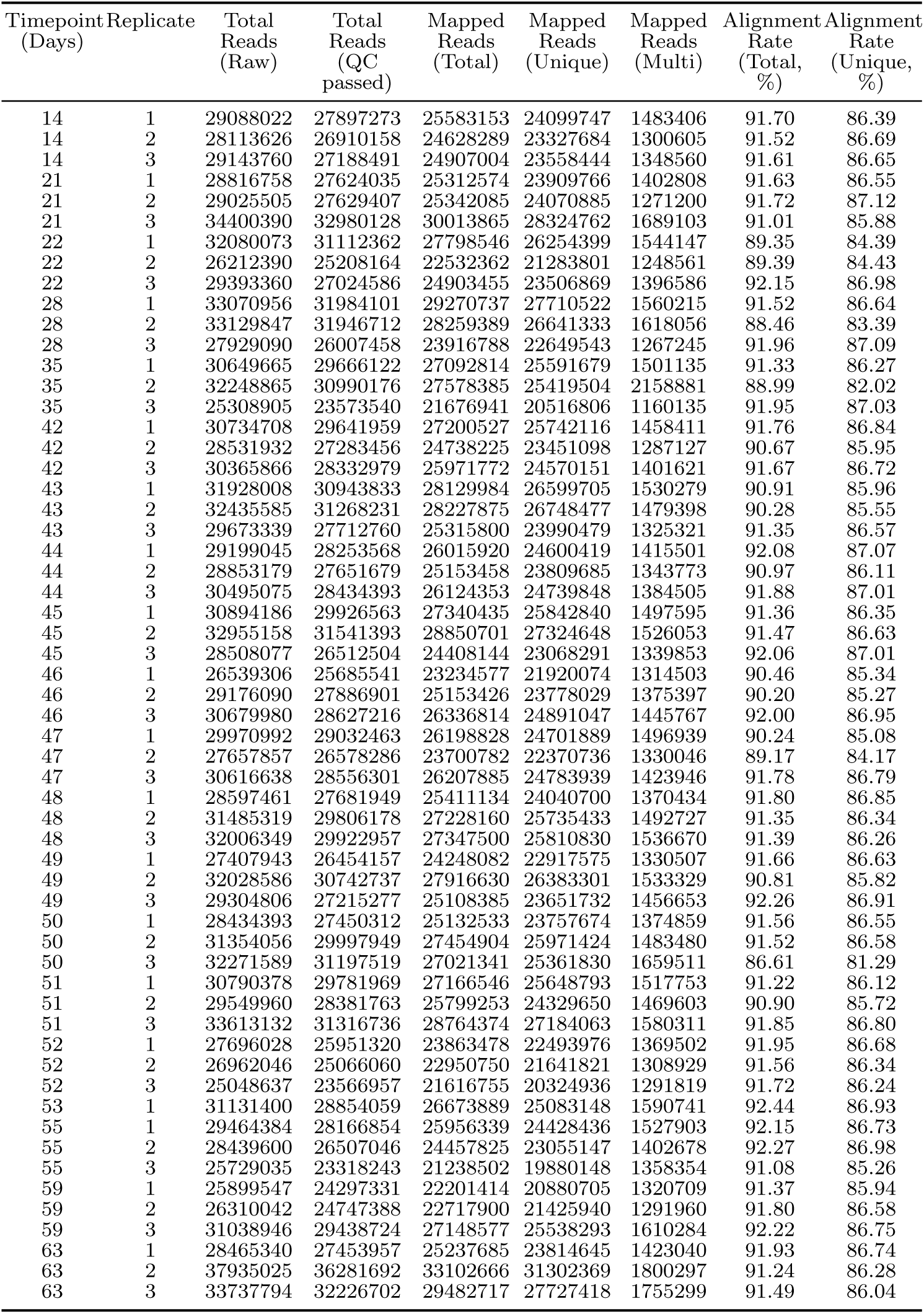
Alignment statistics for *B. napus* cv. ZS11 RNA-seq timeseries. >Samples in ZS11 timeseries, as shown in Figure 1 show consistent alignment rate to the Darmor v10 reference genome [60], with an average of 91.21 %. In every sample, majority of reads are uniquely mapped, with an average of 86.10 %.

| Timepoint<br>(Days) | Replicate | Total<br>Reads<br>(Raw) | Total<br>Reads<br>(QC<br>passed) | Mapped<br>Reads<br>(Total) | Mapped<br>Reads<br>(Unique) | Mapped<br>Reads<br>(Multi) | Alignment<br>Rate<br>(Total,<br>%) | Alignment<br>Rate<br>(Unique,<br>%) |
| --- | --- | --- | --- | --- | --- | --- | --- | --- |
| 14 | 1 | 29088022 | 27897273 | 25583153 | 24099747 | 1483406 | 91.70 | 86.39 |
| 14 | 2 | 28113626 | 26910158 | 24628289 | 23327684 | 1300605 | 91.52 | 86.69 |
| 14 | 3 | 29143760 | 27188491 | 24907004 | 23558444 | 1348560 | 91.61 | 86.65 |
| 21 | 1 | 28816758 | 27624035 | 25312574 | 23909766 | 1402808 | 91.63 | 86.55 |
| 21 | 2 | 29025505 | 27629407 | 25342085 | 24070885 | 1271200 | 91.72 | 87.12 |
| 21 | 3 | 34400390 | 32980128 | 30013865 | 28324762 | 1689103 | 91.01 | 85.88 |
| 22 | 1 | 32080073 | 31112362 | 27798546 | 26254399 | 1544147 | 89.35 | 84.39 |
| 22 | 2 | 26212390 | 25208164 | 22532362 | 21283801 | 1248561 | 89.39 | 84.43 |
| 22 | 3 | 29393360 | 27024586 | 24903455 | 23506869 | 1396586 | 92.15 | 86.98 |
| 28 | 1 | 33070956 | 31984101 | 29270737 | 27710522 | 1560215 | 91.52 | 86.64 |
| 28 | 2 | 33129847 | 31946712 | 28259389 | 26641333 | 1618056 | 88.46 | 83.39 |
| 28 | 3 | 27929090 | 26007458 | 23916788 | 22649543 | 1267245 | 91.96 | 87.09 |
| 35 | 1 | 30649665 | 29666122 | 27092814 | 25591679 | 1501135 | 91.33 | 86.27 |
| 35 | 2 | 32248865 | 30990176 | 27578385 | 25419504 | 2158881 | 88.99 | 82.02 |
| 35 | 3 | 25308905 | 23573540 | 21676941 | 20516806 | 1160135 | 91.95 | 87.03 |
| 42 | 1 | 30734708 | 29641959 | 27200527 | 25742116 | 1458411 | 91.76 | 86.84 |
| 42 | 2 | 28531932 | 27283456 | 24738225 | 23451098 | 1287127 | 90.67 | 85.95 |
| 42 | 3 | 30365866 | 28332979 | 25971772 | 24570151 | 1401621 | 91.67 | 86.72 |
| 43 | 1 | 31928008 | 30943833 | 28129984 | 26599705 | 1530279 | 90.91 | 85.96 |
| 43 | 2 | 32435585 | 31268231 | 28227875 | 26748477 | 1479398 | 90.28 | 85.55 |
| 43 | 3 | 29673339 | 27712760 | 25315800 | 23990479 | 1325321 | 91.35 | 86.57 |
| 44 | 1 | 29199045 | 28253568 | 26015920 | 24600419 | 1415501 | 92.08 | 87.07 |
| 44 | 2 | 28853179 | 27651679 | 25153458 | 23809685 | 1343773 | 90.97 | 86.11 |
| 44 | 3 | 30495075 | 28434393 | 26124353 | 24739848 | 1384505 | 91.88 | 87.01 |
| 45 | 1 | 30894186 | 29926563 | 27340435 | 25842840 | 1497595 | 91.36 | 86.35 |
| 45 | 2 | 32955158 | 31541393 | 28850701 | 27324648 | 1526053 | 91.47 | 86.63 |
| 45 | 3 | 28508077 | 26512504 | 24408144 | 23068291 | 1339853 | 92.06 | 87.01 |
| 46 | 1 | 26539306 | 25685541 | 23234577 | 21920074 | 1314503 | 90.46 | 85.34 |
| 46 | 2 | 29176090 | 27886901 | 25153426 | 23778029 | 1375397 | 90.20 | 85.27 |
| 46 | 3 | 30679980 | 28627216 | 26336814 | 24891047 | 1445767 | 92.00 | 86.95 |
| 47 | 1 | 29970992 | 29032463 | 26198828 | 24701889 | 1496939 | 90.24 | 85.08 |
| 47 | 2 | 27657857 | 26578286 | 23700782 | 22370736 | 1330046 | 89.17 | 84.17 |
| 47 | 3 | 30616638 | 28556301 | 26207885 | 24783939 | 1423946 | 91.78 | 86.79 |
| 48 | 1 | 28597461 | 27681949 | 25411134 | 24040700 | 1370434 | 91.80 | 86.85 |
| 48 | 2 | 31485319 | 29806178 | 27228160 | 25735433 | 1492727 | 91.35 | 86.34 |
| 48 | 3 | 32006349 | 29922957 | 27347500 | 25810830 | 1536670 | 91.39 | 86.26 |
| 49 | 1 | 27407943 | 26454157 | 24248082 | 22917575 | 1330507 | 91.66 | 86.63 |
| 49 | 2 | 32028586 | 30742737 | 27916630 | 26383301 | 1533329 | 90.81 | 85.82 |
| 49 | 3 | 29304806 | 27215277 | 25108385 | 23651732 | 1456653 | 92.26 | 86.91 |
| 50 | 1 | 28434393 | 27450312 | 25132533 | 23757674 | 1374859 | 91.56 | 86.55 |
| 50 | 2 | 31354056 | 29997949 | 27454904 | 25971424 | 1483480 | 91.52 | 86.58 |
| 50 | 3 | 32271589 | 31197519 | 27021341 | 25361830 | 1659511 | 86.61 | 81.29 |
| 51 | 1 | 30790378 | 29781969 | 27166546 | 25648793 | 1517753 | 91.22 | 86.12 |
| 51 | 2 | 29549960 | 28381763 | 25799253 | 24329650 | 1469603 | 90.90 | 85.72 |
| 51 | 3 | 33613132 | 31316736 | 28764374 | 27184063 | 1580311 | 91.85 | 86.80 |
| 52 | 1 | 27696028 | 25951320 | 23863478 | 22493976 | 1369502 | 91.95 | 86.68 |
| 52 | 2 | 26962046 | 25066060 | 22950750 | 21641821 | 1308929 | 91.56 | 86.34 |
| 52 | 3 | 25048637 | 23566957 | 21616755 | 20324936 | 1291819 | 91.72 | 86.24 |
| 53 | 1 | 31131400 | 28854059 | 26673889 | 25083148 | 1590741 | 92.44 | 86.93 |
| 55 | 1 | 29464384 | 28166854 | 25956339 | 24428436 | 1527903 | 92.15 | 86.73 |
| 55 | 2 | 28439600 | 26507046 | 24457825 | 23055147 | 1402678 | 92.27 | 86.98 |
| 55 | 3 | 25729035 | 23318243 | 21238502 | 19880148 | 1358354 | 91.08 | 85.26 |
| 59 | 1 | 25899547 | 24297331 | 22201414 | 20880705 | 1320709 | 91.37 | 85.94 |
| 59 | 2 | 26310042 | 24747388 | 22717900 | 21425940 | 1291960 | 91.80 | 86.58 |
| 59 | 3 | 31038946 | 29438724 | 27148577 | 25538293 | 1610284 | 92.22 | 86.75 |
| 63 | 1 | 28465340 | 27453957 | 25237685 | 23814645 | 1423040 | 91.93 | 86.74 |
| 63 | 2 | 37935025 | 36281692 | 33102666 | 31302369 | 1800297 | 91.24 | 86.28 |
| 63 | 3 | 33737794 | 32226702 | 29482717 | 27727418 | 1755299 | 91.49 | 86.04 |

**Table 3:** Top 25 nodes by degree centrality in the network inferred from *B. napus* cv. Stellar timeseries. *B. napus* gene names are from Darmor v10 reference [60], Arabidopsis gene names are from TAIR 10 [36] and pathways were obtained from FLORID database [9].

| *B. napus* gene | Centrality rank | Arabidopsis id | Symbol | Pathway(s) |
| --- | --- | --- | --- | --- |
| A05p11760.1 | 1 | AT2G33810 | SPL3 | Aging, Photoperiod, Sugar, Hormone |
| A05p34220.1 | 2 | AT3G15270 | SPL5 | Aging, Photoperiod, Sugar, Hormone |
| C05p51970.1 | 3 | AT3G15270 | SPL5 | Aging, Photoperiod, Sugar, Hormone |
| A07p35480.1 | 4 | AT1G69120 | AP1 | Floral integrator |
| A05p05620.1 | 5 | AT2G45660 | SOC1 | Floral integrator |
| A07p31710.1 | 6 | AT1G69120 | AP1 | Floral integrator |
| A10p00850.1 | 7 | AT1G01060 | LHY | Circadian clock |
| C08p32790.1 | 8 | AT3G57390 | AGL18 | Photoperiod |
| C04p06570.1 | 9 | AT2G45660 | SOC1 | Floral integrator |
| A04p26040.1 | 10 | AT2G33810 | SPL3 | Aging, Photoperiod, Sugar, Hormone |
| C07p44700.1 | 11 | AT5G60910 | AGL8 | Floral integrator |
| A03p18910.1 | 12 | AT2G33810 | SPL3 | Aging, Photoperiod, Sugar, Hormone |
| C06p40760.1 | 13 | AT1G69120 | AP1 | Floral integrator |
| A09p47980.1 | 14 | AT3G54990 | SMZ | Aging, Hormone, Photoperiod, Vernalisation |
| A05p10920.1 | 15 | AT2G34720 | NF-YA4 | Photoperiod, Hormone |
| A01p38590.1 | 16 | AT3G15270 | SPL5 | Aging, Photoperiod, Sugar, Hormone |
| C01p05690.1 | 17 | AT4G32980 | ATH1 | Photoperiod |
| C05p00880.1 | 18 | AT1G01060 | LHY | Circadian clock |
| C04p01110.1 | 19 | AT2G46830 | CCA1 | Circadian clock |
| A01p35910.1 | 20 | AT3G18550 | BRC1 | Photoperiod, Meristem identity |
| A05p18570.1 | 21 | AT1G53160 | SPL4 | Aging, Photoperiod, Sugar, Hormone |
| C09p52620.1 | 22 | AT5G60120 | TOE2 | Aging, Photoperiod, Temperature |
| A10p17270.1 | 23 | AT5G60120 | TOE2 | Aging, Photoperiod, Temperature |
| C09p04070.1 | 24 | AT3G28910 | ATMYB30 | Autonomous |
| C04p63380.1 | 25 | AT2G33810 | SPL3 | Aging, Photoperiod, Sugar, Hormone |

**Table 4:** Top 25 nodes by degree centrality in the network inferred from *B. napus* cv. ZS11 timeseries. *B. napus* gene names are from Darmor v10 reference [60], Arabidopsis gene names are from TAIR 10 [36] and pathways were obtained from FLORID database [9].

| *B. napus* gene | Centrality rank | Arabidopsis id | Symbol | Pathway (s) |
| --- | --- | --- | --- | --- |
| C06p29880.1 | 1 | AT1G79430 | APL | Photoperiod |
| A05p16210.1 | 2 | AT5G37260 | RVE2 | Temperature, Circadian clock |
| C02p05760.1 | 3 | AT5G12840 | HAP2A | Photoperiod |
| A04p33940.1 | 4 | AT2G45660 | SOC1 | Floral integrator |
| C04p03360.1 | 5 | AT2G42200 | SPL9 | Hormone, Aging, Sugar |
| A06p47390.1 | 6 | AT5G44190 | GLK2 | Autonomous |
| C02p04280.1 | 7 | AT5G10140 | FLC | Photoperiod, Vernalization |
| A03p25280.1 | 8 | AT2G45660 | SOC1 | Floral integrator |
| C03p70830.1 | 9 | AT3G46640 | PCL1 | Temperature, Circadian clock |
| A02p06580.1 | 10 | AT5G60120 | TOE2 | Temperature, Aging, Photoperiod |
| A10p27400.1 | 11 | AT5G10140 | FLC | Photoperiod, Vernalization |
| A02p41740.1 | 12 | AT5G61850 | LFY | Floral integrator |
| A06p20850.1 | 13 | AT3G46640 | PCL1 | Temperature, Circadian clock |
| A01p26150.1 | 14 | AT3G46640 | PCL1 | Temperature, Circadian clock |
| C04p63380.1 | 15 | AT2G33810 | SPL3 | Hormone, Aging, Photoperiod, Sugar |
| C07p05420.1 | 16 | AT2G17770 | ATBZIP27 | Temperature, Hormone, Aging, Photoperiod, Sugar |
| A09p47980.1 | 17 | AT3G54990 | SMZ | Temperature, Hormone, Aging, Photoperiod, Vernalization |
| A05p18570.1 | 18 | AT1G53160 | SPL4 | Hormone, Aging, Photoperiod, Sugar |
| A09p35240.1 | 19 | AT1G30970 | SUF4 | Vernalisation |
| A03p16730.1 | 20 | AT5G10140 | FLC | Photoperiod, Vernalization |
| A03p04430.1 | 21 | AT5G10140 | FLC | Photoperiod, Vernalization |
| C03p65260.1 | 22 | AT5G61850 | LFY | Floral integrator |
| A05p05620.1 | 23 | AT2G45660 | SOC1 | Floral integrator |
| C03p31440.1 | 24 | AT2G28550 | RAP2.7 | Temperature, Hormone, Aging, Photoperiod, Vernalization |
| C04p69560.1 | 25 | AT2G42200 | SPL9 | Hormone, Aging, Sugar |

**Table 5:**
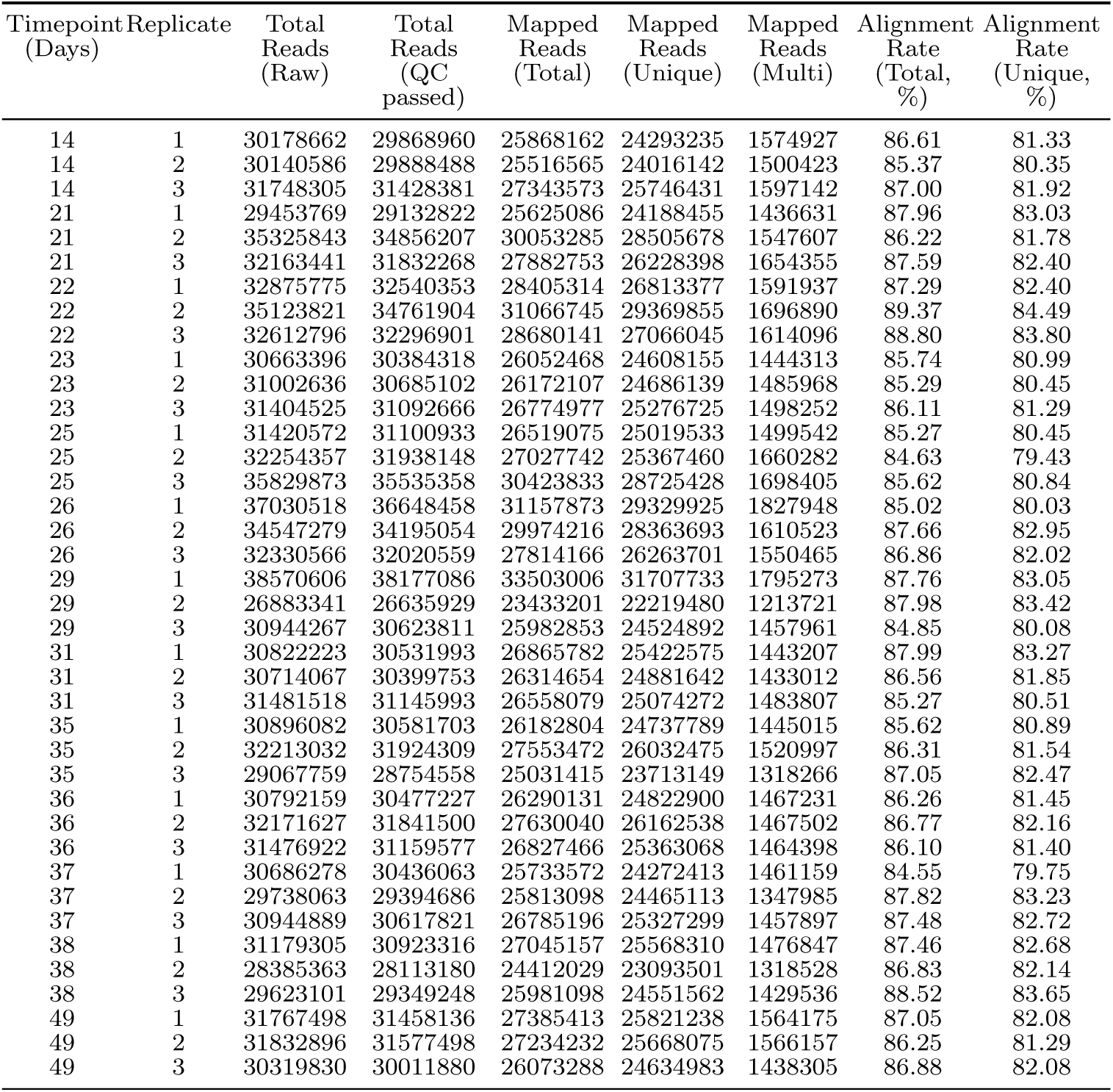
Alignment statistics for *B. napus* cv. Stellar RNA-seq timeseries with short-day, cold treatment. Samples in Stellar timeseries, as shown in Figure 6 (a) show consistent alignment rate to the Darmor v10 reference genome [60], with an average of 86.8 %. In every sample, majority of reads are uniquely mapped, with an average of 81.84 %.

**Table 6:** Identity of protein sequences of *SOC1* orthologues in *B. napus* to Arabidopsis *SOC1*. All *SOC1* proteins in *B. napus* have highly conserved sequences. The MADS domain, the key functional domain consisting of the first 57 amino acids in the Arabidopsis *SOC1* is also highly conserved, except in two orthologues with a substitution from Glycine (G) to Alanine (A) at position 52 from the N-terminal end.

| Orthologue | Identity with Arabidopsis *SOC1* |  |
| --- | --- | --- |
|  | Full Protein | MADS domain |
| A03p25280.1 | 94.39% | 100.0% |
| A05p05620.1 | 95.33% | 98.25% |
| A04p33940.1 | 92.52% | 100.0% |
| C03p30160.1 | 95.33% | 100.0% |
| C04p06570.1 | 95.33% | 98.25% |
| C04p71550.1 | 92.66% | 100.0% |

**Table 7:**
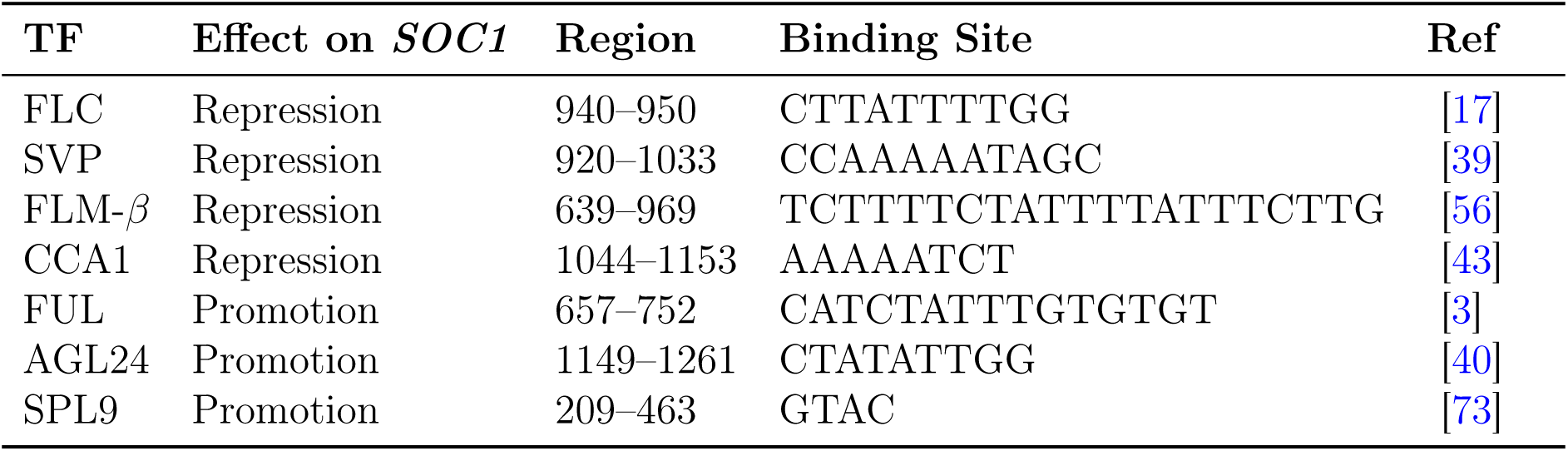
Transcription factors (TFs) regulating *SOC1* expression prior to floral transition through direct binding to the promoter region. TFs that have been experimentally shown to directly bind and regulate *SOC1* expression prior to floral transition and their binding sequences. Binding positions are positions on the negative strand with start codon = 1 in TAIR 10 genome annotation [36]. Modified from the original by Sri *et al*. [68].

**Table 8:** Alignment statistics for *B. napus* cv. Stellar RNA-seq timeseries with short-day and cold treatments. Samples in Stellar timeseries, as shown in Figure 8 alongwith the environmental conditions. The samples are aligned to the Darmor v10 reference genome [60].

| Timepoint<br>(Days) | Replicate | Conditions | Total<br>Reads<br>(Raw) | Total<br>Reads<br>(QC<br>passed) | Mapped<br>Reads<br>(Total) | Mapped<br>Reads<br>(Unique) | Mapped<br>Reads<br>(Multi) | Alignment<br>Rate<br>(Total,<br>%) | Alignment<br>Rate<br>(Unique,<br>%) |
| --- | --- | --- | --- | --- | --- | --- | --- | --- | --- |
| 21 | 1 | 18/15 °C, 16h | 26408972 | 26116762 | 23079155 | 21949803 | 1129352 | 88.37 | 84.04 |
| 21 | 2 | 18/15 °C, 16h | 29257738 | 28948795 | 25650667 | 24320601 | 1330066 | 88.61 | 84.01 |
| 21 | 3 | 18/15 °C, 16h | 28134831 | 27819064 | 24756800 | 23554830 | 1201970 | 88.99 | 84.67 |
| 22 | 1 | 18/15 °C, 8h | 29520012 | 29197554 | 25830916 | 24567306 | 1263610 | 88.47 | 84.14 |
| 22 | 2 | 18/15 °C, 8h | 35112824 | 34689491 | 30760967 | 29285028 | 1475939 | 88.68 | 84.42 |
| 22 | 3 | 18/15 °C, 8h | 35909757 | 35457101 | 31259873 | 29731571 | 1528302 | 88.16 | 83.85 |
| 23 | 1 | 18/15 °C, 8h | 31265438 | 30906338 | 27397303 | 26044327 | 1352976 | 88.65 | 84.27 |
| 23 | 2 | 18/15 °C, 8h | 29887458 | 29557426 | 26244064 | 24791509 | 1452555 | 88.79 | 83.88 |
| 23 | 3 | 18/15 °C, 8h | 31708146 | 31345768 | 27828879 | 26496094 | 1332785 | 88.78 | 84.53 |
| 28 | 1 | 18/15 °C, 8h | 30934706 | 30558054 | 26985907 | 25595606 | 1390301 | 88.31 | 83.76 |
| 28 | 2 | 18/15 °C, 8h | 27929931 | 27564519 | 24481470 | 23202946 | 1278524 | 88.82 | 84.18 |
| 28 | 3 | 18/15 °C, 8h | 30220406 | 29872652 | 26610218 | 25135200 | 1475018 | 89.08 | 84.14 |
| 35 | 1 | 18/15 °C, 8h | 33621146 | 33220961 | 29864743 | 28322503 | 1542240 | 89.90 | 85.25 |
| 35 | 2 | 18/15 °C, 8h | 32976845 | 32689066 | 29469286 | 28001953 | 1467333 | 90.15 | 85.66 |
| 35 | 3 | 18/15 °C, 8h | 31289234 | 30913863 | 27677951 | 26283065 | 1394886 | 89.53 | 85.02 |
| 37 | 1 | 18/15 °C, 16h | 27496683 | 27146294 | 24267295 | 23018286 | 1249009 | 89.39 | 84.79 |
| 37 | 2 | 18/15 °C, 16h | 32331501 | 31935020 | 28734200 | 27271037 | 1463163 | 89.98 | 85.40 |
| 37 | 3 | 18/15 °C, 16h | 37927362 | 37449228 | 33400899 | 31681960 | 1718939 | 89.19 | 84.60 |
| 22 | 1 | 5 °C, 16h | 30739487 | 30410580 | 27053587 | 25755505 | 1298082 | 88.96 | 84.69 |
| 22 | 2 | 5 °C, 16h | 30048219 | 29705879 | 26463637 | 25233533 | 1230104 | 89.09 | 84.94 |
| 22 | 3 | 5 °C, 16h | 35905818 | 35489341 | 31826553 | 30213908 | 1612645 | 89.68 | 85.14 |
| 23 | 1 | 5 °C, 16h | 37898621 | 37472593 | 33146158 | 31521672 | 1624486 | 88.45 | 84.12 |
| 23 | 2 | 5 °C, 16h | 35049527 | 34674034 | 30490549 | 29002808 | 1487741 | 87.93 | 83.64 |
| 23 | 3 | 5 °C, 16h | 33429065 | 33013191 | 27957743 | 26603351 | 1354392 | 84.69 | 80.58 |
| 28 | 1 | 5 °C, 16h | 38291212 | 37834882 | 33087184 | 31334112 | 1753072 | 87.45 | 82.82 |
| 28 | 2 | 5 °C, 16h | 30252551 | 29863873 | 26709085 | 25440420 | 1268665 | 89.44 | 85.19 |
| 28 | 3 | 5 °C, 16h | 30229217 | 29863011 | 26651599 | 25253858 | 1397741 | 89.25 | 84.57 |
| 35 | 1 | 5 °C, 16h | 31769644 | 31364530 | 28194509 | 26811361 | 1383148 | 89.89 | 85.48 |
| 35 | 2 | 5 °C, 16h | 27647080 | 27299647 | 24418218 | 23141758 | 1276460 | 89.45 | 84.77 |
| 35 | 3 | 5 °C, 16h | 30443905 | 29997079 | 26582363 | 25265910 | 1316453 | 88.62 | 84.23 |
| 37 | 1 | 18/15 °C, 16h | 36691375 | 36214062 | 32285550 | 30717887 | 1567663 | 89.15 | 84.82 |
| 37 | 2 | 18/15 °C, 16h | 28559766 | 28211752 | 25464681 | 24179292 | 1285389 | 90.26 | 85.71 |
| 37 | 3 | 18/15 °C, 16h | 27110640 | 26788149 | 24266084 | 23074382 | 1191702 | 90.59 | 86.14 |

